# Laboratory adaptive evolution of thermotolerance is linked to the evolution of a robust proteostasis in S. cerevisiae

**DOI:** 10.1101/2025.01.31.635942

**Authors:** Devi Prasanna Dash, Zainab Zaidi, Akanksha Sharma, Dhwani Dholakia, Manish Rai, Debasish Dash, Kausik Chakraborty

## Abstract

Thermophilic organisms have evolved a proteome that resists thermal denaturation. While the evolution of their complete proteome would require multiple generations, early on, organisms would need to develop strategies to survive at high temperatures despite their thermolabile proteome. We hypothesized that the organisms would do this by reinforcing their proteostasis capacity. We tested this hypothesis using adaptive laboratory evolution of thermotolerance in Saccharomyces cerevisiae and found that the cells reproducibly evolved better proteostasis capacity in short-term evolution experiments. However, rather than improving the global proteostasis capacity, most of the evolved strains demonstrated enhanced capacity to tackle misfolding in the Endoplasmic Reticulum (ER), specifically by increasing their capacity for ER-associated degradation (ERAD). Given the strong selective advantage of these strains, we posit that protein folding in the ER may be exquisitely sensitive to chronic thermal stress and may act as an early indicator for adaptation to higher temperatures.

---

The evolution of thermotolerance provides a fascinating glimpse into the past and the future scope of organisms. Thermotolerant organisms have been harnessed for their thermostable proteome(Haki and Rakshit, 2003; Thomson et al., 2022); they are also crucial in understanding the evolutionary trajectory of organisms (Caspeta et al., 2014; Johnson et al., 2021; Lenski and Bennett, 1993; Riehle et al., 2003; Tenaillon et al., 2012). Understanding their evolutionary trajectory can provide us with early biological warning signs of global warming and identify the fundamental biological processes most vulnerable to increased growth temperatures(Berger et al., 2021; Hoffmann and Sgrò, 2011).

While all biochemical processes are inherently temperature-dependent, many are critically temperature-sensitive. For example, membrane fluidity increases with temperature, increasing the risk of membrane leakage at high temperatures(Fan and Evans, 2015; Hoogerland et al., 2024; Mendoza, 2014); rates of biochemical processes are deregulated at higher temperatures forcing cells to respond to heat shock such that they prevent an uncontrolled increase in reaction rates(Persson et al., 2020). More complex reactions like protein folding exhibit high-temperature sensitivity, especially for soluble, globular proteins that depend upon a hydrophobic effect for folding (Guo et al., 2012; Scalley and Baker, 1997). While many of these temperature-dependent processes could be the subject of selection pressure when an organism evolves thermotolerance, the selection pressure on protein folding has a demonstrable influence and is the most studied aspect in these species(Chakravarty and Varadarajan, 2002, 2000; Das and Gerstein, 2000).

Though the increase in protein stability is a marker of thermotolerance, they are unlikely markers for a short-term increase in growth temperature. Thermotolerant organisms have proteins that are stable at high temperatures and have mutations that make them thermostable(Chakravarty and Varadarajan, 2002, 2000; Das and Gerstein, 2000). Since these mutations have been selected over a long time(Ladenstein and Antranikian, 1998; Steindorff et al., 2024), they are unlikely to be signs of recent selection forces. Logically, cells should evolve a better proteostasis capacity before it has the chance to accumulate mutations throughout their proteome. To have a selective advantage at high temperatures, they should either be able to assist the folding of the existing thermolabile proteome at moderately high temperatures or should be able to take care of toxicity associated with protein misfolding and aggregation. Although this is a logically valid assumption, it has not been tested rigorously in a laboratory setting. This is especially true for eukaryotes, where multiple compartments share the load of folding proteins and ensure the quality of the folded proteins(Lindquist and Kelly, 2011; O’Brien and van Oosten-Hawle, 2016; Powers and Balch, 2013). It is unknown if one or more of the compartments involved in protein folding are particularly vulnerable to elevated temperatures.

Notably, while signatures of selection are of fundamental importance, they would be crucial to identify pathways that can be modified to alter cellular proteostasis. Although we have a comprehensive understanding of protein folding machinery and quality control systems in eukaryotic cells(Hill et al., 2017; Hipp et al., 2019; Jayaraj et al., 2020; Morimoto, 1998), their complex interplay has prevented us from designing pathways or strategies to increase the protein folding efficiency of a cell(Bhattacharya et al., 2020; Calamini et al., 2012; Ghosh et al., 2019; Tokuriki and Tawfik, 2009; VanBogelen et al., 1987). If the short-term evolution experiment changes cellular proteostasis, it can be used to identify critical regulators that reinforce this process.

In this study, we performed a controlled adaptive laboratory evolution (ALE) experiment to generate moderately thermotolerant yeast strains starting from the laboratory strains of S. cerevisiae. We prove the first tenet of short-term evolution: cells evolved better proteostasis with the evolution of thermotolerance. Thermotolerant strains resisted Endoplasmic reticulum protein unfolding stress and not global or cytosolic misfolding, indicating that the ER may be particularly vulnerable to elevated temperatures. These strains upregulated the ER-associated degradation pathway, which seems to be the conserved signature of selection for thermostability in S. cerevisiae. Few of these strains also showed better protein folding capability. Thus, reinforcement of different branches of proteostasis may allow cells to survive better at elevated temperatures. Importantly, we found that although single-gene knockouts can faithfully recapitulate thermotolerance, better proteostasis cannot be. This suggests that altered proteostasis is a more complex multigenic trait that follows the evolution of thermotolerance in laboratory environments.

## Results

### Adaptive Laboratory evolution to generate thermotolerance in S. cerevisiae

To recreate an early step in the evolution of thermotolerance, we chose the mesophilic S. cerevisiae as the model eukaryote and performed adaptive laboratory evolution (ALE) at 40°C (ten degrees higher than the physiological growth temperature used in the laboratory) by repeated dilution and regrowth (Figure 1A). While a single replicate of ALE would provide one route to achieving thermotolerance, they would not reveal the preference of routes. Additionally, many of the mutations in the genome may be associated with the trait either causally (mutations that increase thermotolerance) or because they co-occurred with a causal mutation. We used parallel replicates of ALE to identify the preferred routes to achieve intermediate thermotolerance (Tenaillon et al., 2012). This also decreased the noise in identifying causal mutations: genes or gene modules mutated in multiple strains are more likely to be linked causally to the selected trait. To circumvent the problem of cross-well contamination and contamination from external sources, we evolved two genetically different strains of S. cerevisiae (BY4741 and S288C) (BY: URA-/MATa and S288C: URA+/MATα), interspersed in the plate (Figure 1A). Additional controls were put in place to ensure (cross)contamination-free evolution of the strains (please refer to Material and Methods for details)(Figure S1A, S1B, S1C, and S1D). Following ALE for ∼600 generations, single colonies were isolated from each of the wells and further processed. The thermotolerant BY4741-derived strains are prefixed with TT with a number unique to each strain (the strain names and the background of the parental strains are provided in Supplemental Table S1) The analysis provided in the work focuses exclusively on TT strains on the background of BY4741, unless otherwise stated explicitly.

**Figure 1:**
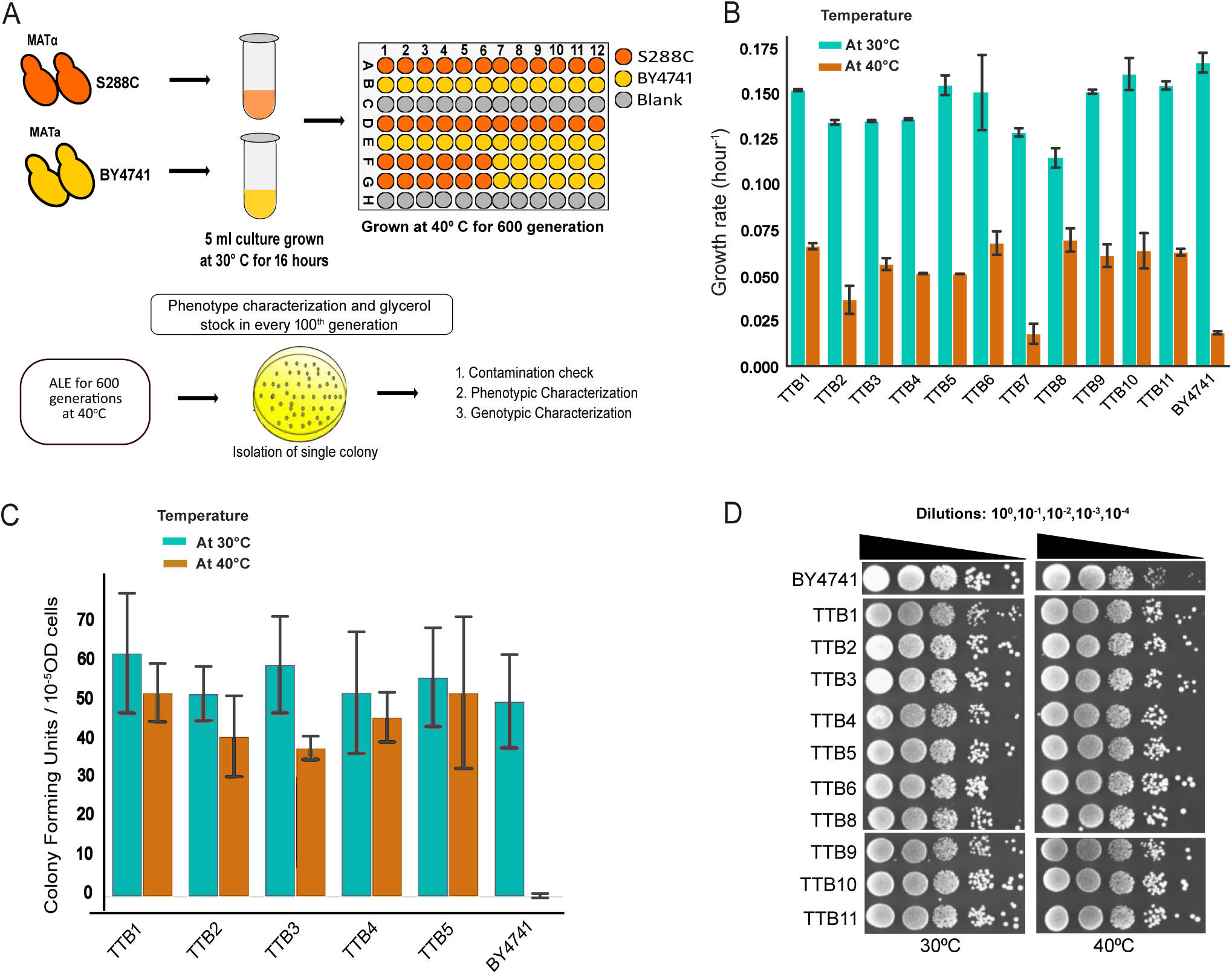
S. cerevisiae strains are adapted to higher temperatures during the process of ALE at 40°C for 600 generation. (A) Schematic overview of the Adaptive Laboratory Evolution (ALE) approach at 40°C using deep multi-well plates. Yeast strains BY4741 and S288C were subjected to high-temperature conditions (40°C) in nutrient-rich YPD broth. Cells were cultured to saturation and re-inoculated into fresh media, with this cycle repeated for approximately 600 generations to promote adaptation. Following this evolution phase, single colonies were isolated from each independent population, and both phenotypic and genotypic characterizations were performed to assess adaptive changes. (B) Growth rates of representative thermoevolved BY4741 strains (TTB1 to TTB11) and the unevolved parent strain BY4741 in YPD medium at 30°C and 40°C. (C) Cell viability of thermos-evolved TT strains compared to the unevolved BY4741 control strain at 30°C and 40°C, measured via colony-forming units (CFUs). (D) Drop dilution assay of thermoevolved TT strains alongside the unevolved BY4741 control, using 10-fold serial dilutions up to 10^−4^ on YPD agar plates at 30°C and 40°C.

To confirm the genetic fixation of thermotolerance, we made stocks of the TT strains, grew them at 30^0^C for a few generations, and then cultured them back at 40^0^C. Growth rates of most of the representative TT strains, when grown continuously at 40^0^C, were higher than the parental BY4741 (referred to as BY hereafter) (Figure 1B, S1E). The strains however showed a marginal but non-significant growth defect at 30^0^C (Figure 1B) indicating a slight, if any, change in the optimal temperature of growth. Similar results were obtained for the competitive fitness experiments reported in an accompanying manuscript (Zaidi et al., 2024).

The growth rate of the evolved strains could be higher than the parental strain at 40^0^C through either of the following strategies: an increase in the proliferation rate at higher temperatures or a decrease in cell death at higher temperatures. Survival assay shows that the evolved strains have lower cell death at higher temperatures (Figure 1C) indicating that the evolved strains manage to find solutions to toxicity associated with high-temperature growth. Drop dilution assays on solid media also confirmed that the TT strains had evolved an advantage to grow at higher temperatures (Figure 1D). Thus, using the ALE strategy we evolved strains that are better at handling chronic thermal stress than the parental BY.

### Genome sequencing reveals routes to thermotolerance

Of the isolated colonies, we obtained quality genome sequencing data from 29 thermotolerant strains derived from BY4741, and 19 thermotolerant strains from S288C. 367 unique mutations were detected in the BY4741-derived strains, while 144 unique mutations were obtained in the S288c-derived strains. These mutations comprised both nonsense and missense mutations (Table S2). We focused on genes with at least one non-sense mutation in one of the thermotolerant strains. Since, premature stop codons terminated these genes, deleting these genes would likely show a thermotolerant phenotype if these mutations imparted thermotolerance. Assuringly, two of these genes that accumulated stop codons during ALE, lrg1 (Jarolim et al., 2013) and mrn1 (Huang et al., 2018), are already known to confer thermotolerance to BY4741 when deleted, validating the approach and the methodology of ALE. We validated the rest of the genes using the deletion collection of S. cerevisiae (Winzeler et al., 1999). All the single gene deletion strains (*inh1Δ, vip1Δ, seg2Δ, kex2Δ, mot3Δ,* and *swa2Δ*) were competitively fitter than BY at 40^0^C (Figure 2A). Their fitness at 30^0^C was lower than BY(Figure 2B), recapitulating the trend shown by TTB strains. This was further confirmed by growth on solid media (Figure S2A, B). To rule out artifacts of suppressor mutations, we reconstituted the deletion strains and validated the results (Figure S2C, D). Thus, we could recapitulate two known genes and identify six new genes whose deletions increase thermotolerance in the background of BY4741.

**Figure 2:**
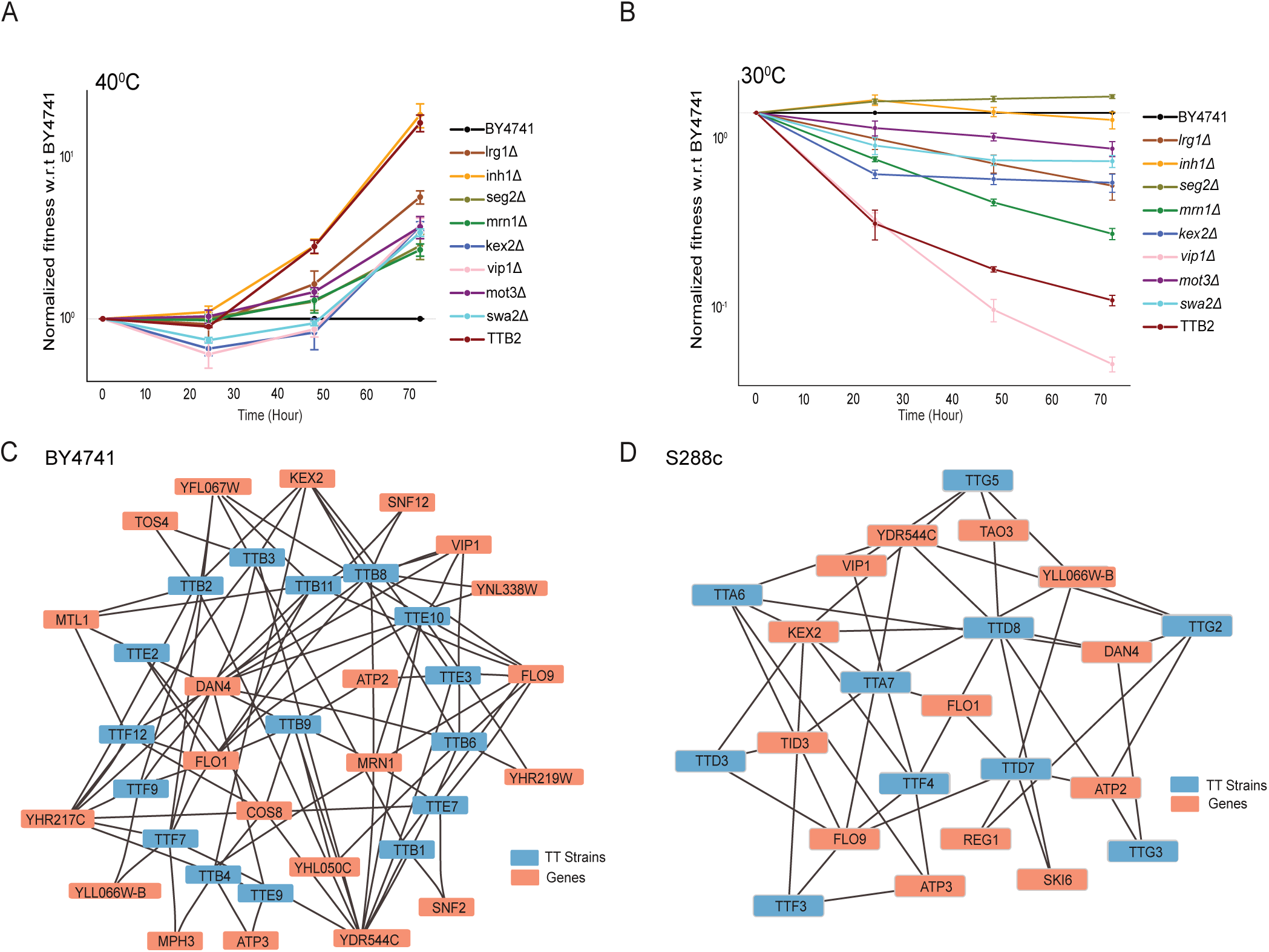
Genotype characterization from whole genome sequencing reveals routes to thermotolerance. (A) Competitive fitness of selected strains from the deletion library in the BY4741 background at 40°C, normalized to the unevolved BY4741 strain at 40°C, grown in YPD media. (B) Competitive fitness of selected strains from the deletion library in the BY4741 background at 40°C, normalized to the unevolved BY4741 strain at 30°C, grown in YPD media. (C) Network of TT strains generated from the parental BY4741 strain, displaying clusters of mutations in similar genes. Clustering is based on the Jaccard metric, illustrating associations between TT strains and their respective mutated genes. (D) Mutations belonging to similar genes in many TT strains generated from parental S288C were clustered base Jaccard metric. The TT strains associated with genes are shown in the network.

Genes that accumulated missense mutations in multiple strains were clustered based on the Jacard metric(Chung et al., 2019)(Figure 2C, D, and S2E); there was no definite pattern in the pathways that accumulated mutations. However, we found the disomy of Chromosome III to be a recurrent feature in both TTB (6 out of 29) and TTS (3 out of 19) (TableS3 and S2F). Trisomy of ChrIII in the background of diploid S. cerevisiae is known to impart thermotolerance(Yona et al., 2012). Our study recapitulated that an increase in the dosage of ChrIII can increase the thermotolerance of BY. In addition, we found the loss of mitochondrial DNA to be a recurrent event in the process of evolution, which is the focus of discussion in an accompanying manuscript (Zaidi et al., 2024).

To check if the disomy of ChrIII was sufficient to confer thermotolerance in the background of the haploid BY strain, we used the Δecm1, which is known to lead to ChrIII disomy BY4741(Hughes et al., 2000). Δecm1 was more thermotolerant than BY (Figure S2A, D), indicating that the disomy is sufficient to confer thermotolerance.

Taken together, genome sequencing indicated two sufficient routes to thermotolerance: loss of function of genes whose deletion confers thermotolerance and the duplication of the third chromosome. The third route involving mitochondrial chromosome loss is insufficient to confer thermotolerance. It is a more complex phenomenon linked to proteostasis and is the subject of an accompanying manuscript (Zaidi et al., 2024).

### Thermotolerant strains evolve better tolerance to ER proteotoxicity

We next asked if the short-term evolution of thermotolerance fixed mutations that altered cellular proteostasis. Most experiments are performed at 30°C unless otherwise stated to check this fixation. Since decreasing translation is a common strategy to increase proteostasis capacity during misfolding stresses (Clay et al., 2023; Steffen et al., 2012), we asked if the evolved cells may use such a strategy to improve their proteostasis capacity. We checked for translation using a GFP expressed under a constitutive promoter in the evolved strains. There was no significant difference in single-cell fluorescence of GFP(Figure S3A). Since the half-life of GFP in BY is exceptionally long, the protein production rates had to be roughly similar in the thermotolerant and the parental BY at 30°C. Thus, protein biogenesis does not differ in the evolved strains.

The other way to increase proteostasis capacity would be to better assist the folding of proteins. To check the folding arm of proteostasis, we used the folding sensors of NAT-R (Nourseothricin Acetyl Transferase)(Ghosh et al., 2019). The protein in its active form can confer tolerance to the antibiotic Nourseothricin (Nat). Cells that fold the folding sensor well would tolerate higher concentrations of Nat without altering their ability to tolerate Nat. Out of the strains tested, TTB2 and TTB3, two stains that harbored ChrIII disomy, were able to fold the folding-sensor TS22 better than BY at 30°C (Figure 3A) and 34°C (Figure 3B, S3B). However, another evolved line, TTB1 did not exhibit any increase in TS22 activity. This showed that at a physiological temperature or a mild heat shock condition (34°C), TTB2 and TTB3 could fold TS22 better than the parental BY or TTB1 that lacked ChrIII disomy. This indicated that the strains harboring ChrIII disomy may be associated with an increased ability to fold proteins. However, given that TTB1 doesn’t show an increase in the cytosolic protein folding capacity, this was not a general feature of the evolved strains.

**Figure 3:**
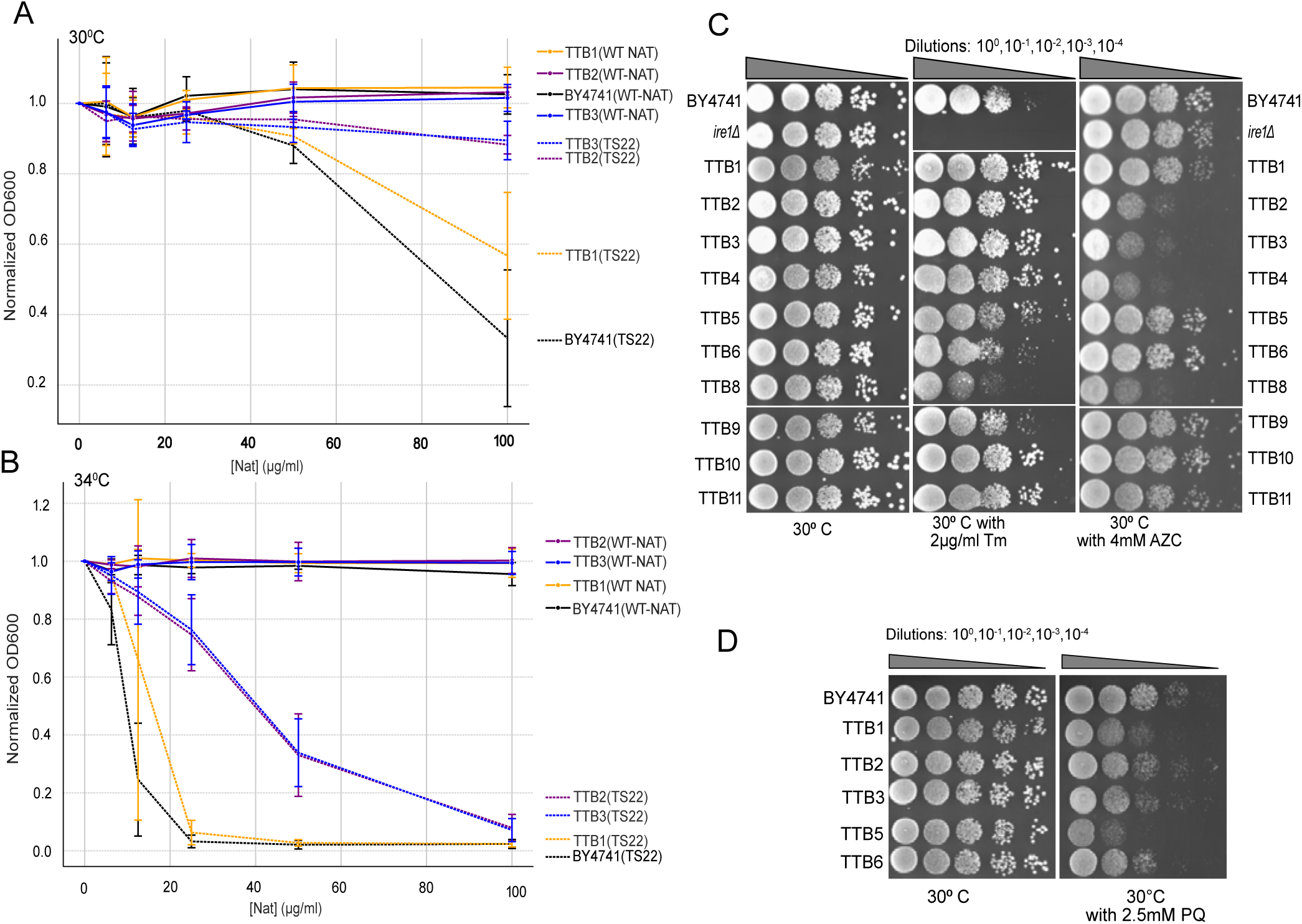
Thermotolerant strains develop a better tolerance to ER proteotoxicity. (A) MIC assay conducted with varying concentrations of Nourseothricin (ClonNAT) at 30°C in YPD medium. OD at 600 nm was measured and normalized to assess inhibitory effects across different concentrations. (B) MIC assay conducted with varying concentrations of Nourseothricin (ClonNAT) at 34°C in YPD medium. OD at 600 nm was measured and normalized to assess inhibitory effects across different concentrations. (C) Drop dilution assay of TT strains in the YPD agar plate incubated at 30°C in presence of Tunicamycin(Tm; 2µg/ml), Azetidine-2-carboxylic acid (AZC; 2.5mM) with unevolved BY4741 and ⊗ire1 as control. (D) Drop dilution assay of TT strains in the YPD agar plate incubated at 30°C in presence of Paraquat(PQ; 2.5mM) with unevolved BY4741 as control.

Cells may improve proteostasis capacity by increasing their tolerance to protein misfolding. To examine if the evolved strains were adept at handling stress associated with misfolding, we monitored their fitness in the presence of small molecules that induce misfolding. Specifically, we used Azetidine-2-carboxylic acid (AZC), which misincorporates in place of proline to induce misfolding in all compartments (global misfolding), and Tunicamycin (Tm), which inhibits protein glycosylation in the endoplasmic reticulum (ER) and triggers ER-specific misfolding. Ten representative TT strains were exposed to these proteotoxic stresses at 30°C to assess whether their evolved mutations helped stabilize proteostasis at physiological temperatures. At 30°C, several strains (TTB1, 2, 3, 4, 5, 10, 11) displayed greater tolerance to Tm than the BY strain, while others exhibited hypersensitivity (TTB8) or similar sensitivity (TTB6, 9). Notably, tolerance to Tm increased further at 40°C (Figure S3C), suggesting that most strains evolved to genetically stabilize ER proteostasis under proteotoxic stress. In contrast, none of the strains demonstrated enhanced tolerance to AZC at 30°C (Figure 3C) or 40°C (Figure S3C), with the exception of TTB9 and 11. Many strains showed increased sensitivity to AZC (TTB2, 3, 4, 8), and strains with ChrIII disomy were particularly sensitive. These results indicate that the evolution of thermotolerance in these strains was accompanied by increased resistance to ER-specific misfolding stress but did not extend to a broader resistance against global proteotoxic stress.

Many of these insults to global proteostasis or chronic growth at high temperatures are known to switch on an environmental stress response (ESR) pathway (Gasch et al., 2000) driven primarily by oxidative stress response genes. These genes also play a critical role in tolerating ER stress(Maity et al., 2016). We asked if ER stress tolerance or thermotolerance in these strains was due to their enhanced resistance to oxidative stress. We challenged the cells with paraquat (Pq), which induces oxidative stress. None of the strains tested (except TTB2) showed higher resistance to Pq at 30°C (Figure 3D). The same was true at 40°C (Figure S3D), suggesting that the evolution of thermotolerance or ER stress tolerance in the evolved lines did not require better ESR or tolerance to oxidative stress.

Thus, some of the thermotolerant strains that harbored ChrIII disomy evolved and enhanced the ability to fold folding-compromised mutant proteins. Additionally, most strains evolved an increased ability to handle proteotoxicity due to ER-associated misfolding. However, none seem to have evolved an ability to handle global proteotoxicity more efficiently than the parental BY strain.

### Single gene deletions partially recapitulate proteostasis rewiring

To investigate the individual contributions of specific mutations and ChrIII disomy to proteostasis rewiring, we used representative deletion strains. These deletions either 1) resulted in a stop codon mutation in the thermotolerant strains, which, when deleted in the BY strain, conferred thermotolerance (Figure 2A, B)) or 2) induced ChrIII disomy (e.g., ecm1).

Deletion of inh1, vip1, or ecm1 increased thermotolerance but did not increase the foldability of TS22 either at 30°C (Figure 4A, B) or 34°C (Figure 4C). This demonstrated that enhanced thermotolerance does not guarantee better foldability of the folding sensor. This also suggests that, though strains with ChrIII disomy exhibit better folding capacity, the disomy alone (in Δecm1) is insufficient to ensure better protein folding capacity.

**Figure 4:**
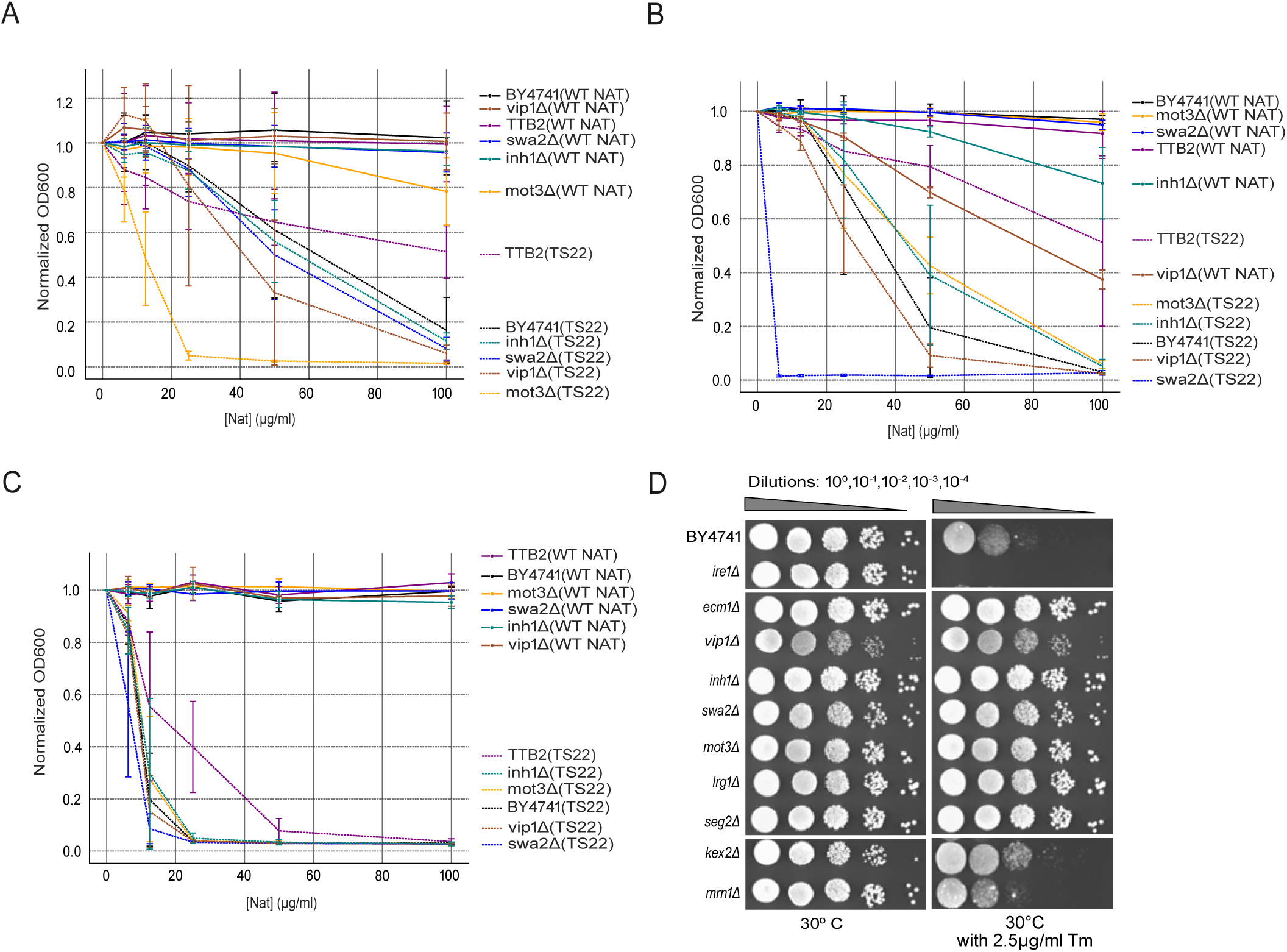
Rewiring of proteostasis is partially recapitulated by single gene deletions. (A and B) MIC assay for reconstructed deletion strains conducted with varying concentrations of Nourseothricin (ClonNAT) at 30°C in YPD medium. OD at 600 nm was measured and normalized to assess inhibitory effects across different concentrations. (C) MIC assay for reconstructed deletion strains conducted with varying concentrations of Nourseothricin (ClonNAT) at 34°C in YPD medium. OD at 600 nm was measured and normalized to assess inhibitory effects across different concentrations. (D) Drop dilution assay of reconstructed deletion strains in the YPD agar plate incubated at 30°C in presence of Tunicamycin(Tm; 2µg/ml) with unevolved BY4741 and Δire1 as control.

In stark contrast to protein folding, single deletion of multiple genes (ecm1, inh1, mot3, seg2 and mrn1) conferred better ER stress tolerance (Figure 4D and Figure 4SA). Surprisingly, an overlapping but distinct set of single gene deletions (inh1, vip1, swa2) conferred fitness in AZC(Figure 4SB). This suggests that while certain gene deletions enhance thermotolerance and resistance to proteotoxic stress, they do so without impacting the protein-folding aspect of proteostasis. Notably, despite these deletions conferring fitness advantages with AZC exposure, the evolved strains themselves did not show improved fitness in AZC relative to the parental BY strain.

### Ire1-independent ER stress response is activated in the TTB strains

To check for the possible mechanisms through which TTB strains tolerate ER stress, we deleted ire1 in the background of these strains. Ire1 deletion confers extreme Tm-sensitivity to BY; even in this background, representative evolved strains were more resistant to Tm than BY (Figure 5A). Thus, TTB strains could counter ER stress without the involvement of Ire1-dependent signaling of UPR^ER^. Additionally, we confirmed that the thermotolerance of the TTB lines was independent of Ire1 (Figure 5B, C).

**Figure 5:**
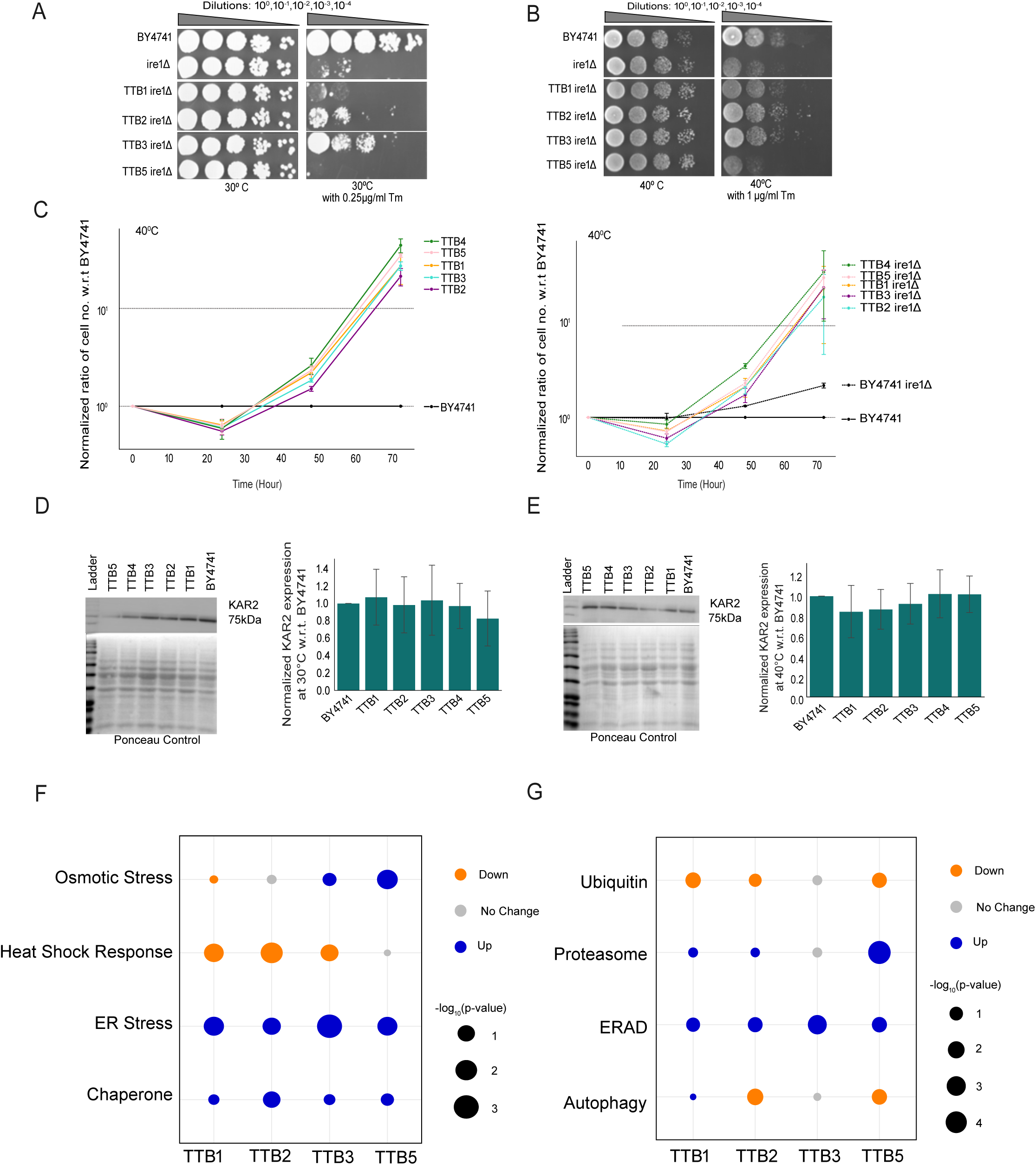
Thermotolerance is independent of UPRER stress response pathway and Ire1-independent ER stress response is activated in the TTB strains. (A) Drop dilution assay for log phase cultures in YPD agar plate in presence of Tunicamycin(Tm; 0.25µg/ml) incubated at 30°C with unevolved BY4741 and Δire1 in BY4741 as control. (B) Drop dilution assay for log phase cultures in YPD agar plate in presence of Tunicamycin(Tm; 0.25µg/ml) incubated at 40°C with unevolved BY4741 and Δire1 in BY4741 as control. (C) Relative fitness of TTB strains at 40°C (left panel). Relative fitness of TTB:ire1Δ strains at 40°C (right panel). Normalization done w.r.t. unevolved parental BY4741 strain. (D) Western blot showing KAR2 expression in strains grown at 30°C. 20μg of total protein from whole cell lysate was loaded and probed with anti-KAR2. Normalization done w.r.t. KAR2 expression in unevolved parental BY4741 strain. (E) Western blot showing KAR2 expression in strains grown at 40°C. 20μg of total protein from whole cell lysate was loaded and probed with anti-KAR2. Normalization done w.r.t. KAR2 expression in unevolved parental BY4741 strain. (F) Bubble Plot of Stress Response Pathways Across Different TT Strains Based on Transcriptomic Data. The bubble plot illustrates the relationship between fold change and p-values for the expression of stress response pathways for four different TT strains (TTB1, TTB2, TTB3, TTB5). The colors of the bubbles represent the log2 fold changes of gene expression, with blue indicating upregulation and orange indicating downregulation. The overlaid circle size in each cell reflects the significance of the change, represented by the -log_10_ of the p-value. Larger circles indicate more significant changes (lower p-values), while smaller circles indicate less significant changes (higher p-values). Stress response pathways are listed on the y-axis, and the strains are represented on the x-axis. (G) Bubble Plot of Degradation Pathways Across Different TTB Strains Based on Transcriptomic Data. Degradation pathways are listed on the y-axis, and the strains are represented on the x-axis.

Since Kar2 is exquisitely sensitive to Ire1 activation, we checked if Ire1 signaling was upregulated in the evolved strain using Kar2 levels as an indicator. In the absence of any ER stressor, Kar2 levels did not increase significantly in the TT strains over BY4741, indicating that the evolved lines did not upregulate the Ire1 pathway in basal state at 30°C as well as 40°C (Figure 5D, E); this supports the idea that the TT lines evolved ER-stress tolerance independent of the Ire1 pathway.

Gene set enrichment analysis of the transcriptomic data of TTB strains at 30°C indicated that the ER stress response pathway genes(Travers et al., 2000) were upregulated in all four representative strains (Figure 5F and S5B). Interestingly, the heat shock response pathway was downregulated in these strains (Figure 5F), indicating that ER-stress resistance is independent of the heat shock response.

Since these evolved lines tolerate proteotoxicity better than BY and we hypothesized that they have better clearance mechanisms, we check for gene sets that comprise the genes involved in different protein degradation pathways. Corroborating the ER-stress tolerance data, ERAD genes(Travers et al., 2000) were upregulated in most mutants (Figure 5G and S5A).

Taken together, this suggests that the evolution of thermotolerance is linked to better tolerance towards ER-proteotoxicity. The strains reinforce ER proteostasis by upregulating ER-stress response and ERAD pathways independent of the canonical Ire1 and heat-shock response pathways.

## Discussion

Here, we link the laboratory evolution of thermotolerance to the evolution of better proteostasis in haploid test strains of S. cerevisiae. Using ALE, we generated S. cerevisiae lines that were thermotolerant and obtained crucial information on proteostasis modulation. We learned that thermotolerance was better linked to resistance to ER-associated proteotoxicity than to global protein misfolding stress. The evolved strains handled the stress independently of the canonical Ire1 pathway or the oxidative quality control pathway, two pathways known to control ER proteostasis(Maity et al., 2016b; Ron and Walter, 2007). Notably, a subset of the strains that contained ChrIII disomy was better at folding mutant proteins. However, none of the single gene deletions could recapitulate this in the background of BY.

### Proteostasis is reinforced during the short-term evolution of thermotolerance

While long-term evolution at high temperatures selects for a thermostable proteome, a typical mesophilic organism has a large part of the proteome that aggregates at high temperatures (Jarzab et al., 2020). Hence, it is highly unlikely that the organism can stabilize all the thermolabile proteins in a short time. There is an orthogonal route that the cells may take to assist folding of the thermolabile proteome. Cells have mechanisms to assist the folding of unstable proteins (Domnauer et al., 2021; Mogk et al., 2018); there are specialized proteins (molecular chaperones) that help proteins fold (Kim et al., 2013; Saibil, 2013) along with metabolites that stabilize the transition states (chemical chaperones) to assist folding (Bolen, 2004; Dandage et al., 2015; Ignatova and Gierasch, 2006; Taneja and Ahmad, 1994). Increasing the abundance of these machinery is likely a way to solve the problem of temperature stress in the short term while allowing time for the organism to accumulate mutations to stabilize the proteome in the long run. We show here that the thermolabile proteome is partially stabilized in some of the short-term lab-evolved thermotolerant strains of S. cerevisiae. Strains with improved folding efficiency also showed increased sensitivity to misfolding stress induced by AZC, suggesting that while these strains evolved an enhanced capacity for protein folding, they were less capable of clearing terminally misfolded proteins. Notably, this enhanced folding capacity could not be attributed to elevated molecular chaperone expression levels (Figure 5F). This concurs well with previous reports that did not find the overexpression of chaperones in evolved thermotolerant lines of E. coli (Tenaillon et al., 2012). We speculate that changes in metabolite levels may support this enhanced folding by acting as chemical chaperones to help bypass kinetic traps that become more prevalent at higher temperatures. However, we have yet to identify any specific metabolites with significant alterations.

### Selection pressure arises mostly from compromised ER protein folding

Given the preponderance of ER stress resistance in the strains tested, it is tempting to speculate that the primary selection pressure during chronic high-temperature growth acts on the foldability of proteins in the ER. ER is involved in two unique post-translational modifications that chaperone folding: glycosylation and disulfide bond formation(Breitling and Aebi, 2013; Sevier and Kaiser, 2002). Both of these modifications are important for folding ER-targeted proteins, as disruption of either leads to the induction of UPR^ER^(Ferreira et al., 2017). Since disulfide bonds can reduce the entropy of the unfolded chains(Matsumura et al., 1989; Pantoliano et al., 1987), ER proteins that depend upon disulfides for folding may be sensitive to conditions that increase the entropy of the unfolded state. Folding at high temperatures is known to be plagued by an increase in entropy of the unfolded protein chain and, hence, may render the ER proteins more vulnerable to misfolding. Since the thermotolerant strains overexpress the ERAD pathway genes(Chen et al., 1988; Travers et al., 2000), they are better capable of handling misfolding due to either thermal stress or due to Tm-induced misfolding of glycosylation-dependent proteins. However, the evolved strains did not depend on Ire1 for their enhanced thermotolerance or increased ability to handle ER stress. This suggests that other pathways can regulate the expression of ERAD genes.

Surprisingly, thermotolerant strains did not evolve resistance to global misfolding stress as monitored by AZC tolerance. A few single gene deletions showed resistance, but the evolved strain did not. While this suggests that cytosolic misfolding may not be a significant hurdle to yeast at the moderately high temperatures used in this study (40°C), this also indicates that cytosolic misfolding may be managed well by the various co-translational folding apparatus present in the cytosol(Kim et al., 2013; Nissley et al., 2022; Ruan et al., 2017).

### Limitations

The nuclear genomes of the strains contained multiple changes, and these altered the transcriptomes differently in each of the strains, having convergent and divergent trends. The converging trends allow us to obtain system-level information regarding pathways that may be critical for proteostasis. However, epistasis and other interactions prevent us from pinpointing any single gene or process that is critical for altering proteotoxicity or proteostasis. However, given the system’s level changes, we would need new models and methods for understanding the evolution of these complex traits.

### Outlook

Our work shows that thermotolerance can evolve in the absence of a better protein folding arm or better tolerance to global proteotoxic stress, as seen from the single gene deletions that are thermotolerant but do not have better proteostasis. The evolution of a better proteostasis is more complex and evolves after a driver mutation increases the thermotolerance of the organism. Our study would be critical in delving into two aspects of cellular proteostasis: the connection between ER stress and tolerance to proteotoxicity and the ability to increase the folding capacity of cells through cellular engineering. The evolved strains show that it is possible to design cells with better protein folding capacity. Although we have not pinpointed pathways that increase the folding capacity, our report has the potential to form the basis of such studies.

## Acknowledgment

We acknowledge Shikha Rao and Soumen Kundu for critically reviewing and editing the manuscript. KC acknowledges CSIR FIRST funding MLP2303 for funding the study and CSIR Institute of Genomics and Integrative Biology (CSIR-IGIB) for its HPC and Highthroughput sequencing facility. KC is an India Alliance Senior Fellow. Devi Prasanna Dash (DPD) acknowledges DBT; Z.Z acknowledges CSIR-IGIB; A.S acknowledges UGC; Dhwani Dholakia acknowledges DBT for fellowship support.

## References

1. Berger D, Stångberg J, Baur J, Walters RJ. 2021. Elevated temperature increases genome-wide selection on de novo mutations. Proceedings of the Royal Society B 288:20203094.

2. Bhattacharya K, Weidenauer L, Luengo TM, Pieters EC, Echeverría PC, Bernasconi L, Wider D, Sadian Y, Koopman MB, Villemin M. 2020. The Hsp70-Hsp90 co-chaperone Hop/Stip1 shifts the proteostatic balance from folding towards degradation. Nat Commun 11:5975.

3. Bolen DW. 2004. Effects of naturally occurring osmolytes on protein stability and solubility: issues important in protein crystallization. Methods 34:312–322.

4. Breitling J, Aebi M. 2013. N-linked protein glycosylation in the endoplasmic reticulum. Cold Spring Harb Perspect Biol 5:a013359.

5. Calamini B, Silva MC, Madoux F, Hutt DM, Khanna S, Chalfant MA, Saldanha SA, Hodder P, Tait BD, Garza D. 2012. Small-molecule proteostasis regulators for protein conformational diseases. Nat Chem Biol 8:185–196.

6. Caspeta L, Chen Y, Ghiaci P, Feizi A, Buskov S, Hallström BM, Petranovic D, Nielsen J. 2014. Altered sterol composition renders yeast thermotolerant. Science (1979) **346**:75–78.

7. Chakravarty S, Varadarajan R. 2002. Elucidation of factors responsible for enhanced thermal stability of proteins: a structural genomics based study. Biochemistry 41:8152–8161.

8. Chakravarty S, Varadarajan R. 2000. Elucidation of determinants of protein stability through genome sequence analysis. FEBS Lett 470:65–69.

9. Chen C, Bonifacino JS, Yuan LC, Klausner RD. 1988. Selective degradation of T cell antigen receptor chains retained in a pre-Golgi compartment. J Cell Biol 107:2149–2161.

10. Chung NC, Miasojedow B, Startek M, Gambin A. 2019. Jaccard/Tanimoto similarity test and estimation methods for biological presence-absence data. BMC Bioinformatics 20:644.

11. Clay KJ, Yang Y, Clark C, Petrascheck M. 2023. Proteostasis is differentially modulated by inhibition of translation initiation or elongation. Elife 12:e76465.

12. Dandage R, Bandyopadhyay A, Jayaraj GG, Saxena K, Dalal V, Das A, Chakraborty K. 2015. Classification of chemical chaperones based on their effect on protein folding landscapes. ACS Chem Biol 10:813–820.

13. Das R, Gerstein M. 2000. The stability of thermophilic proteins: a study based on comprehensive genome comparison. Funct Integr Genomics 1:76–88.

14. Domnauer M, Zheng F, Li L, Zhang Yanxiao, Chang CE, Unruh JR, Conkright-Fincham J, McCroskey S, Florens L, Zhang Ying. 2021. Proteome plasticity in response to persistent environmental change. Mol Cell 81:3294–3309.

15. Fan W, Evans RM. 2015. Turning up the heat on membrane fluidity. Cell 161:962–963.

16. Ferreira RB, Wang M, Law ME, Davis BJ, Bartley AN, Higgins PJ, Kilberg MS, Santostefano KE, Terada N, Heldermon CD. 2017. Disulfide bond disrupting agents activate the unfolded protein response in EGFR-and HER2-positive breast tumor cells. Oncotarget 8:28971.

17. Gasch AP, Spellman PT, Kao CM, Carmel-Harel O, Eisen MB, Storz G, Botstein D, Brown PO. 2000. Genomic expression programs in the response of yeast cells to environmental changes. Mol Biol Cell 11:4241–4257.

18. Ghosh A, Gangadharan A, Verma M, Das S, Matai L, Dash DP, Dash D, Mapa K, Chakraborty K. 2019. Cellular responses to proteostasis perturbations reveal non-optimal feedback in chaperone networks. Cellular and Molecular Life Sciences 76:1605–1621.

19. Guo M, Xu Y, Gruebele M. 2012. Temperature dependence of protein folding kinetics in living cells. Proceedings of the National Academy of Sciences 109:17863–17867.

20. Haki GD, Rakshit SK. 2003. Developments in industrially important thermostable enzymes: a review. Bioresour Technol 89:17–34.

21. Hill SM, Hanzén S, Nyström T. 2017. Restricted access: spatial sequestration of damaged proteins during stress and aging. EMBO Rep 18:377–391.

22. Hipp MS, Kasturi P, Hartl FU. 2019. The proteostasis network and its decline in ageing. Nat Rev Mol Cell Biol 20:421–435.

23. Hoffmann AA, Sgrò CM. 2011. Climate change and evolutionary adaptation. Nature 470:479–485.

24. Hoogerland L, van den Berg SPH, Suo Y, Moriuchi YW, Zoumaro-Djayoon A, Geurken E, Yang F, Bruggeman F, Burkart MD, Bokinsky G. 2024. A temperature-sensitive metabolic valve and a transcriptional feedback loop drive rapid homeoviscous adaptation in Escherichia coli. Nat Commun 15:9386.

25. Huang C-J, Lu M-Y, Chang Y-W, Li W-H. 2018. Experimental evolution of yeast for high-temperature tolerance. Mol Biol Evol 35:1823–1839.

26. Hughes TR, Roberts CJ, Dai H, Jones AR, Meyer MR, Slade D, Burchard J, Dow S, Ward TR, Kidd MJ. 2000. Widespread aneuploidy revealed by DNA microarray expression profiling. Nat Genet 25:333–337.

27. Ignatova Z, Gierasch LM. 2006. Inhibition of protein aggregation in vitro and in vivo by a natural osmoprotectant. Proceedings of the National Academy of Sciences 103:13357–13361.

28. Jarolim S, Ayer A, Pillay B, Gee AC, Phrakaysone A, Perrone GG, Breitenbach M, Dawes IW. 2013. Saccharomyces cerevisiae genes involved in survival of heat shock. G3: Genes, Genomes, Genetics **3**:2321–2333.

29. Jarzab A, Kurzawa N, Hopf T, Moerch M, Zecha J, Leijten N, Bian Y, Musiol E, Maschberger M, Stoehr G. 2020. Meltome atlas—thermal proteome stability across the tree of life. Nat Methods 17:495–503.

30. Jayaraj GG, Hipp MS, Hartl FU. 2020. Functional modules of the proteostasis network. Cold Spring Harb Perspect Biol 12:a033951.

31. Johnson MS, Gopalakrishnan S, Goyal J, Dillingham ME, Bakerlee CW, Humphrey PT, Jagdish T, Jerison ER, Kosheleva K, Lawrence KR. 2021. Phenotypic and molecular evolution across 10,000 generations in laboratory budding yeast populations. Elife 10:e63910.

32. Kim YE, Hipp MS, Bracher A, Hayer-Hartl M, Ulrich Hartl F. 2013. Molecular chaperone functions in protein folding and proteostasis. Annu Rev Biochem 82:323–355.

33. Ladenstein R, Antranikian G. 1998. Proteins from hyperthermophiles: stability and enzymatic catalysis close to the boiling point of water. Biotechnology of Extremophiles 37–85.

34. Lenski RE, Bennett AF. 1993. Evolutionary response of Escherichia coli to thermal stress. Am Nat 142:S47–S64.

35. Lindquist SL, Kelly JW. 2011. Chemical and biological approaches for adapting proteostasis to ameliorate protein misfolding and aggregation diseases–progress and prognosis. Cold Spring Harb Perspect Biol 3:a004507.

36. Maity S, Rajkumar A, Matai L, Bhat A, Ghosh A, Agam G, Kaur S, Bhatt NR, Mukhopadhyay A, Sengupta S. 2016a. Oxidative homeostasis regulates the response to reductive endoplasmic reticulum stress through translation control. Cell Rep 16:851–865.

37. Matsumura M, Becktel WJ, Levitt M, Matthews BW. 1989. Stabilization of phage T4 lysozyme by engineered disulfide bonds. Proceedings of the National Academy of Sciences 86:6562–6566.

38. Mendoza D de. 2014. Temperature sensing by membranes. Annu Rev Microbiol 68:101–116.

39. Mogk A, Bukau B, Kampinga HH. 2018. Cellular handling of protein aggregates by disaggregation machines. Mol Cell 69:214–226.

40. Morimoto RI. 1998. Regulation of the heat shock transcriptional response: cross talk between a family of heat shock factors, molecular chaperones, and negative regulators. Genes Dev 12:3788–3796.

41. Nissley DA, Jiang Y, Trovato F, Sitarik I, Narayan KB, To P, Xia Y, Fried SD, O’Brien EP. 2022. Universal protein misfolding intermediates can bypass the proteostasis network and remain soluble and less functional. Nat Commun 13:3081.

42. O’Brien D, van Oosten-Hawle P. 2016. Regulation of cell-non-autonomous proteostasis in metazoans. Essays Biochem 60:133–142.

43. Pantoliano MW, Ladner RC, Bryan PN, Rollence ML, Wood JF, Poulos TL. 1987. Protein engineering of subtilisin BPN’: enhanced stabilization through the introduction of two cysteines to form a disulfide bond. Biochemistry 26:2077–2082.

44. Persson LB, Ambati VS, Brandman O. 2020. Cellular control of viscosity counters changes in temperature and energy availability. Cell 183:1572–1585.

45. Powers ET, Balch WE. 2013. Diversity in the origins of proteostasis networks—a driver for protein function in evolution. Nat Rev Mol Cell Biol 14:237–248.

46. Riehle MM, Bennett AF, Lenski RE, Long AD. 2003. Evolutionary changes in heat-inducible gene expression in lines of Escherichia coli adapted to high temperature. Physiol Genomics 14:47–58.

47. Ron D, Walter P. 2007. Signal integration in the endoplasmic reticulum unfolded protein response. Nat Rev Mol Cell Biol 8:519–529.

48. Ruan L, Zhou C, Jin E, Kucharavy A, Zhang Y, Wen Z, Florens L, Li R. 2017. Cytosolic proteostasis through importing of misfolded proteins into mitochondria. Nature 543:443–446.

49. Saibil H. 2013. Chaperone machines for protein folding, unfolding and disaggregation. Nat Rev Mol Cell Biol 14:630–642.

50. Scalley ML, Baker D. 1997. Protein folding kinetics exhibit an Arrhenius temperature dependence when corrected for the temperature dependence of protein stability. Proceedings of the National Academy of Sciences 94:10636–10640.

51. Sevier CS, Kaiser CA. 2002. Formation and transfer of disulphide bonds in living cells. Nat Rev Mol Cell Biol 3:836–847.

52. Steffen KK, McCormick MA, Pham KM, MacKay VL, Delaney JR, Murakami CJ, Kaeberlein M, Kennedy BK. 2012. Ribosome deficiency protects against ER stress in Saccharomyces cerevisiae. Genetics 191:107–118.

53. Steindorff AS, Aguilar-Pontes MV, Robinson AJ, Andreopoulos B, LaButti K, Kuo A, Mondo S, Riley R, Otillar R, Haridas S. 2024. Comparative genomic analysis of thermophilic fungi reveals convergent evolutionary adaptations and gene losses. Commun Biol 7:1124.

54. Taneja S, Ahmad F. 1994. Increased thermal stability of proteins in the presence of amino acids. Biochemical Journal 303:147–153.

55. Tenaillon O, Rodríguez-Verdugo A, Gaut RL, McDonald P, Bennett AF, Long AD, Gaut BS. 2012. The molecular diversity of adaptive convergence. Science (1979) **335**:457–461.

56. Thomson RES, Carrera-Pacheco SE, Gillam EMJ. 2022. Engineering functional thermostable proteins using ancestral sequence reconstruction. Journal of Biological Chemistry 298.

57. Tokuriki N, Tawfik DS. 2009. Chaperonin overexpression promotes genetic variation and enzyme evolution. Nature 459:668–673.

58. Travers KJ, Patil CK, Wodicka L, Lockhart DJ, Weissman JS, Walter P. 2000a. Functional and genomic analyses reveal an essential coordination between the unfolded protein response and ER-associated degradation. Cell 101:249–258.

59. VanBogelen RA, Acton MA, Neidhardt FC. 1987. Induction of the heat shock regulon does not produce thermotolerance in Escherichia coli. Genes Dev 1:525–531.

60. Winzeler EA, Shoemaker DD, Astromoff A, Liang H, Anderson K, Andre B, Bangham R, Benito R, Boeke JD, Bussey H. 1999. Functional characterization of the S. cerevisiae genome by gene deletion and parallel analysis. Science (1979) **285**:901–906.

61. Yona AH, Manor YS, Herbst RH, Romano GH, Mitchell A, Kupiec M, Pilpel Y, Dahan O. 2012. Chromosomal duplication is a transient evolutionary solution to stress. Proceedings of the National Academy of Sciences 109:21010–21015.

62. Zaidi Z, Dash DP, Sharma A, Kundu S, Bhatt S, Rao S, Padia K, Rai M, Chakraborty K. 2024. Fitness defects due to cytosolic protein misfolding in S. cerevisiae can be alleviated by decreasing mitochondrial protein import capacity. *bioRxiv* 2024–2028.

