## Supplemental files for "Laboratory adaptive evolution of thermotolerance is linked to the evolution of a robust proteostasis in S. cerevisiae"

### Supplemental methods

#### Yeast strains, plasmids, and growth conditions

The Saccharomyces cerevisiae strains used in this study were cultivated on rich YPD medium, which contains 1% (w/v) yeast extract, 2% (w/v) peptone, and 2% (w/v) dextrose. Details about the strains of Saccharomyces cerevisiae used in this study can be found in the key resource table. The selection process for deletion strains and plasmid-transformed strains was carried out using artificial drop-out media. Unless otherwise noted, cultures were typically incubated at 30°C with 200 rpm of shaking Different strains used for the study were inoculated in YPD at 0.1/0.2 O.D600 from an overnight primary culture and grown to mid-log phase before harvesting for further experiments.

#### Yeast gene Knockout and gene integration:

WT NAT/variant NAT i.e.TS22 gene integration was carried out in the ura3 disrupted locus of TT strains and BY4741. For generating specific gene knockout strains, complete ORF was replaced by the KanMX4 gene. Homologous recombination was used for all gene integration and knockout procedures. Details of the primers used are given in the key resource table.

#### Generation of thermotolerant Saccharomyces cerevisiae strains through Adaptive Laboratory Evolution (ALE)

For Adaptive Laboratory Evolution (ALE), Saccharomyces cerevisiae strains of different mating types, specifically BY4741 *(MATa his3Δ1 leu2Δ0 lys2Δ0 ura3Δ0*) and S288C (*MATα SUC2 gal2 mal2 mel flo1 flo8-1 hap1 ho bio1 bio6*) (obtained from the American Type Culture Collection (ATCC)), were used as the parent strain for the adaptive evolution experiment (ALE). Single colonies from each parental strain were inoculated into 5 ml of YPD broth and were grown overnight to saturation at 30°C. Saturated primary cultures were re-inoculated at an initial O.D.600 of 0.1 into 400μl of YPD broth in 2.2 ml deep multi-well plates. The growth conditions for the secondary cultures were then shifted directly to 40°C in a water bath with shaking at 100 rpm. For ALE, a total of 72 parental strains (36 BY4741 strains and 36 S288C strains) were considered. The cells were cultured for 600 generations at 40°C. Passages were performed every 16-20 hours, transferring 1% of the inoculum from each well to 400 μl of fresh media in new deep well plates. Glycerol stocks were stored at -80°C every 100 generations, and the growth phenotypes and contaminations of the evolved strains were monitored.

**Ruling out contamination:**

Ruling out contamination was performed for TT strains every 100 generations, as well as for single colonies isolated from TT strains after 600 generations.

**Contamination by other microorganisms:** By using both microscopy and streaking in LB and YPD agar plates, contamination with different microorganisms, such as bacteria and other yeasts, was checked. Crystal violet stain from the Gram staining kit (K001-1KT) from Himedia and Leica DMI6000 microscope were used for this purpose. Contamination by other organisms was ruled out based on colony morphology and growth in YPD and LB plates.

**Cross-contamination between two mating type strains:** Cross-contamination was checked for TT strains from the BY4741 background by simply spotting the cells on an SD-ura agar plate. BY4741one of the parent strain used for ALE, is uracil auxotroph and cannot grow on synthetic defined media deficient in uracil but S288C on the other hand can grow on SD-ura plate. Possible cross-contamination in the case of TT strains from S288C background was ruled out by performing a mating-type PCR (Huxley et al., 1990). The specific PCR amplification product for the parental BY4741 (MATa) strain is 492 bp, and for the S288C (MATα) strainis 369 bp. This helps rule out cross-contamination from S288C background strains.

**Growth assay:**

The primary cultures were grown overnight for approximately 18 hours at 30°C in deep multi-well plates. The saturated cultures were re-inoculated in fresh YPD medium at an OD600 of approximately 0.1 in a honeycomb plate for growth assay in the Bioscreen instrument. Growth assays for all TT strains were performed at 30°C for 24 hours. For the growth assay at 40°C, primary cultures were preconditioned at 40°C for 24 hours, followed by growth at 40°C for 48 hours using the Bioscreen system. The growth curves of each evolved strain were plotted and compared with the unevolved parental strains. In-house Python scripts were used to plot the growth curves, and growth rate calculations were performed using the Grofit model in R (Kahm et al., 2010).

**Cell Viability of TT Strains at 30°C and 40°C:**

Primary cultures were grown at 30ºC and 40ºC (With preconditioning at 40ºC for 24 hours) for 24 hours. Subsequently, 100µl of 1 OD cells diluted to 1:10,000 were plated in YPD Agar and allowed to grow at 30ºC for 48 hours. After 48 hours, CFUs of evolved strains were calculated and compared with unevolved strain.

**Drop dilution assay:**

From saturated primary cultures, secondary cultures were inoculated at 0.1 OD and grown at 30°C. After growing secondary cultures to 1 OD, they were serially diluted 10-fC1 from 0 to $10^{-4}$. Following this, 5µl of every dilution was spotted onto YPD agar plates, and the cells were incubated under different growth conditions.

**Competitive Fitness assay:**

An equal number of cells evolved as well as unevolved GFP tagged strains in a 1:1 ratio were taken and grown at 30°C for 72 hours. Every 24 hours, the ratio between no of evolved to unevolved cells was measured in flow cytometry. The log ratio of evolved to unevolved cells, normalized to the unevolved strain, was plotted to compare the competitive fitness of evolved TT strains relative to unevolved parental strains under different growth conditions.

**Genome sequencing and analysis**:

The glass bead method was used to isolate the S.cerevisae genomic DNA. Before preparing the libraries, the genomic DNA quality and concentration were assessed by using an agarose gel and a Qubit high-sensitivity DNA kit, respectively. The Illumina Nextera XT library preparation kit (15031942) was utilized for the preparation of the library, and indexing was carried out using the dual indexing kit (Nextera XT Index kit v2). For library quality as well as quantity check DNA HS bioanalyzer kit (5067-4626) and Qubit HS DNA kit (Q32854) were used respectively. The pooled gDNA libraries were sequenced on the Illumina Hiseq 2500 platform with 300*2 bp paired-end reads,with a target coverage of approximately 40X per sample. FastQ files were generated by Illumina bcl2 fastq Conversion software. FastQC was used to assess the quality of the reads and low-quality reads were filtered by trimmomatic(Bolger et al., 2014). Read alignment to reference S. cerevisiae genome was done by BWA(Li and Durbin, 2010). For post-processing of BAM files and removal of duplicate reads, SAMtools was used (Li et al., 2009). GATK was used for variant calling (McKenna et al., 2010).

**RNA sequencing and analysis:**

Saturated primary culture of yeast strains i.e. TTB1, TTB2, TTB3, TTB5, ecm1Δ, inh1Δ re-inoculated in fresh YPD media at 0.1 OD, and cells were grown up to 0.5 OD. AT 0.5 OD, cells were harvested and mRNA was isolated and enriched using the Dynabeads™ mRNA Purification Kit (Catalog number: 61006).RNA sequencing library preparation was done by using the Ion Total RNA-Seq Kit v2 (Catalog number: 4479789). Utilizing the mentioned protocol, sequencing was performed on PI™ Hi-Q™ Sequencing 200 Kit (Catalog number: A26433) on the Ion Proton™ System. After sequencing, FastQC (Babraham Bioinformatics) was used to check the FASTQ files quality (Bolger et al., 2014) and Trimmomatic was used to eliminate low-quality reads from the FASTQ files. Hisat2 was used to align the sequenced reads to the yeast genome (Kim et al., 2019).Cuffdiff was used to achieve differential expression of genes (Trapnell et al., 2013). Using the R Wilcox package, the Mann-Whitney-U test was employed for pathway-specific analysis.

**Western Blotting:**

0.1 OD and ~0.35 OD cells from overnight grown saturated primary culture were re-inoculated in fresh YPD and were grown at 30°C and 40°C respectively for 6 hours. Then cultures were harvested and proteins were isolated by alkaline lysis method and the concentrations of proteins were estimated by PierceTM BCA kit (Thermo Scientific, Cat no-23225). For SDS-PAGE and western blotting,30ug of total protein from each sample were loaded in 10% SDS-PAGE gel and the proteins in the gel were transferred to methanol-activated PVDF membrane (Millipore; ISEQ00010). After the transfer, the membrane was stained with ponceau stain, and the captured image was used as loading control. After blocking the membrane for 90 minutes with 2% BSA in TBST, primary antibodies for KAR2 were used for the western blotting experiment. The blots were incubated at 4C overnight and after overnight incubation, blots were washed with TBST thrice for 15 mins. The membrane was then incubated with an HRP-conjugated secondary antibody (anti-rabbit) for two hours at room temperature. After the secondary antibody incubation, the membranes were washed thrice for 15 minutes each to get rid of non-specific binding of antibodies. Secondary antibodies conjugated with horse radish peroxidase (HRP) were used and the SYNGENE gel documentation system was used to capture the developed blot results. Immobilon Western Chemiluminescent HRP substrate (Millipore, WBKLS0500) was used to develop the blots. ImageJ software (Schneider et al., 2012) was utilized for protein quantification, and Ponceau images were used for loading control for normalization.

**MIC of TT strains:**

From the overnight primary culture (400ul of culture in a deep multiwell plate for 18 hours), 0.1 OD of the 400ul culture was grown for 12-16 hours at 30°C, in a deep multiwell plate with a concentration gradient clonnat. The concentration gradient for Minimum Inhibitory Concentration (MIC) ranges from 0 to 100 ug/ml for clonnat. Two-fC1 serial dilution of concentration gradient was considered. Growth readings were obtained using a multi-plate reader in TECAN. A similar protocol was followed for Tunicamycin and AZC with concentration gradients ranging from 0 to 5 µg/ml and 0 to 10 mM, respectively.

**Quantification And Statistical Analysis**

Student t-test used for statistical analysis. The Wilcox package in R was utilized to perform a pathway-specific analysis, utilizing the Mann-Whitney-U test.

**Data And Software Availability**

All data is provided in the manuscript. The genome sequencing files can be found under bioproject accession PRJNA1126056.

### List of chemicals Used

| Chemicals | Source | Catalogue number |
|---|---|---|
| 0.5M EDTA PH 8.0-2X500ML. | Himedia | ML014 |
| 1M Tris-Cl pH 7.4 | Himedia | ML028-500ML |
| 2-mercaptoethanol | MP Biomedicals | 219024283 |
| 2-Propanol | Sigma | I9516-500ML |

| | | |
|---|---|---|
| Acetic acid glacial, AR | Himedia | AS119-2.5L |
| Adenine hemisulfate | Sigma | A9126-100G |
| Adenine hemisulphate salt | Sigma | A9126-100G |
| Agar powder bacteriological grade | Himedia | GRM026 |
| Agarose special, Low EEO for Mol Bio | Himedia | MB002-500G |
| Ammonium chloride | MP Biomedicals | 219462390 |
| Ammonium persulfate | Sigma | A3678 |
| Anti-Mouse IgG (whole molecule)–Peroxidase antibody produced in rabbit | Sigma | A9044-2ML |
| Bovine serum albumin fraction-V | Himedia | GRM105-100G |
| Calcium Chloride 2H2O @12%-500G | Himedia | MB034 |
| Casein(Tryptone) | Himedia | RM014-500G |
| Chemiluminescent HRP substrate millipore | Millipore | WBKLS0500 |
| COOMASSIE* BRILLIANT | MP Biomedicals | 219068225 |
| Cycloheximide | Sigma | 01810-5G |
| Deoxyribonucleic acid, single stranded from salmon testes | Sigma | D9156-5X1ML |
| D-Glucose | MP Biomedicals | 219402491 |
| Dimethyl sulfoxide, for molecular biology | Sigma | D8418-100ml |
| Dithiothreitol | Sigma | 43816-50ml |
| DNA Gel Loading Dye (6x) | Thermo Fisher Scientific | |
| EDTA | Sigma | E6758 |
| Ethanol | Millipore | 1009832500 |
| Ethidium bromide(EtBr) | Sigma | E7637 |
| G 418 DISULFATE SALT | Sigma | A1720-25G |
| Glass beads, acid washed 425-600 µm (30-40 U.S. sieve) | Sigma | G8772-500G |
| Glycerol | MP Biomedicals | 4800689 |
| Glycerol Ultrapure | Sigma | G2025-500mL |
| Glycine | MP Biomedicals | 0219482591 |
| Glycine, molecular biology reagent | MP Biomedicals | 219482594 |
| Grams Stain-Kit | Himedia | K001 |
| Hydrochloric Acid | Himedia | AS004-2.5L |
| Immobilon-PSQ PVDF Membrane | Millipore | ISEQ00010 |
| Isoamyl alcohol | Millipore | 1009791000 |
| L Histidine | Sigma | H6034-100G |
| L- lysine monohydrate | Sigma | L5626-100G |
| L -Methionine | Sigma | M9625-100G |
| L-Alanine | Sigma | A7627-100G |
| L-Arginine | Sigma | A5006-500G |
| L-Asparagine | Sigma | A4159-100G |

| | | |
|---|---|---|
| L-Aspartic acid sodium salt | Sigma | 11189-100G |
| L-Azetidine-2-carboxylic acid | Sigma | A0760-1G |
| L-Cysteine hydrochloride monohydrate | Sigma | |
| L-Glutamic acid sodium salt | Sigma | G3407-100G |
| L-Glutamine | Sigma | G8540-100G |
| Liqui-Gel™ acrylamide and bisacrylamide mixture (29:1) | MP Biomedicals | 4800803 |
| Lithium acetate dihydrate | Sigma | 62393 |
| L-Leucine | Sigma | L8000-100G |
| L-Lysine dihydrochloride | Sigma | L5626-500G |
| L-phenylalanine | Sigma | P2126-100G |
| L-Proline | Sigma | P0380-100G |
| L-Serine | Sigma | S4500-100G |
| L-Threonine | Sigma | T8625-100G |
| L-Threonine | Sigma | T8625-100G |
| L-Tryptophan | Sigma | T0254-100G |
| L-Tryptophane | Sigma | T0254-100G |
| L-Tyrosine | Sigma | T3754-100G |
| L-Tyrosine | Sigma | T3754-100G |
| Luria Bertani Agar, Miller | Himedia | M1151-2.5KG |
| Luria Bertani Broth, Miller | Himedia | M1245-2.5KG |
| L-Valine | Sigma | V0500-100G |
| Magnesium chloride hexahydrate | Himedia | MB040-500G |
| Manganese (II) chloride tetrahydrate, Extra pure | Himedia | GRM685-500G |
| Methanol-AR grade | Himedia | AS059-2.5L |
| Millex-GP sterile syringe filter PES 0.22um 33mm | Millipore | SLGPR33RS |
| Millex-HP 33mm 0.45um PES sterile syringe filter | Millipore | SLHPR33RB |
| Nourseothricin(clonNAT) | Jena bioscience | AB-102XL |
| Nuclease free water | Ambion | AM9937 |
| Page ruler prestained protein ladder | Thermo Fisher Scientific | 26620 |
| Paraquat (Methyl viologen dichloride hydrate 98%) | Sigma | 856177 |
| PBS (Phosphate Buffered Saline | Himedia | TS1101 |
| PEG, Poly(ethylene glycol) 3350 | Sigma | 202444 |
| Peptone | Himedia | RM001 |
| Phenol:Chloroform:Isoamyl Alcohol 25:24:1,pH 4.5 | Thermo Fisher Scientific | AM9720 |
| PIPES(Piperazine-1,4-bis(2-ethanesulfonic acid)) | Sigma | P6757-100G |
| Ponceau S Solution | Sigma | P7170-1L |
| RNAase A | Sigma | 10109169001 |
| SDS | Sigma | L3771 |

| | | |
|---|---|---|
| Sealing Film, Sterile, 50 Sheets | BR Biochem Life Sciences | BC0418-S |
| SOB medium (Hanahan's Broth) | Himedia | M1252 |
| SOC Growth Medium | Himedia | G015-500G |
| Sodium chloride | MP Biomedicals | 219484801 |
| Sodium hydroxide | Himedia | GRM467 |
| TEMED | Sigma | T9281 |
| TRIS base | MP Biomedicals | 210313301 |
| Triton X-100 | Sigma | 93443-500ML |
| Trizol | Thermo Fisher Scientific | 15596018 |
| Tunicamycin | Sigma | T7765-10mg |
| TWEEN 20 | Sigma | P1379-1L |
| Uracil | Sigma | U1128-100G |
| Yeast Extract | Himedia | RM027 |
| Yeast Nitrogen Base w/o amino acids | Himedia | M151 |
| YPD (YEPD) Growth Agar | Himedia | G038-500G |
| YPD Broth- 500GM | Himedia | M1363-500G |
| NTC or ClonNAT | Jena Bioscience | AB102XL |

### Kits and Reagents and Software Used

#### I. List of Antibodies Used

| | | |
|---|---|---|
| Kar2 (polyclonal) | Anti-Rabbit (Santa Cruz Biotechnology) | sc-33630 |
| Anti-Rabbit secondary | Santa Cruz Biotechnology | SC2030 |

#### II. Oligonucleotides

| REAGENT | SOURCE |
|---|---|
| **Primer for integration of NAT variants fused with DsRed at C terminal construct along with URA**<br>**NAT_ORF_DSRed_integration_For_primer**<br>GGATAAGTTTTGACCATCAAAGAAGGTTAATGTGGCTGTGGAGGTCGACGGTATCGATAAGCTTC<br>**NAT_ORF_DSRed_integration_Rev_primer**<br>CATGAAGCTTTTTCTTTCCAATTTTTTTTTTTTCGTCATTGAGAGTGCACCACGCTTTTCAATTC<br>**Confirmation primer for NAT cassette integration**<br>**1. 5 prime confirmation**<br>**NAT_int_5Prime_check_FP** ATTGAGGCTACTGCGCCAAT<br>**NAT_int_5Prime_check_RP** GTGCGGCCATCAAAATGTAT<br>**2. 3 prime confirmation**<br>**NAT_int_3prime_check_FP** CTCCAGTAATTCCTTGGTGG<br>**NAT_int_3prime_check_RP** CGTCCATCTTTACAGTCCTG<br><br>**3. Checking NAT ORF**<br>**NAT_ORF_amplify_FP** CAGTTCTCACATCACATCCG | This Study |

| | |
|---|---|
| **NAT_ORF_amplify_RP** ACCAGCACCTGCTCCGAATT | |
| **inh1_for_int_devi**<br>CGCATTACTACAGCACACTTTTATACAGTTCCACAATAGAATATGGAAGCTTCGTACGCTG CAGGTC<br>**inh1_rev_int_devi**<br>TTCTAAAAAAAAAAAAAAAAAAGCTTCTGCGGAAACGCATGATTAGCATAGGCCACTAGT GGATCTGA<br>**confirmation_inh1**<br>**5 prime confirmation**<br>**CTAACTCGCTATAGCCTTTTCAGTG**<br>**3 prime confirmation**<br>**TTGATTTTTATTCCAACAAGAAGGT** | This Study |
| **Final for deletion vip1**<br>**vip1_for_int_devi**<br>TTCAAAAGCATCTCGTAGCATATTAATATATTGCAGAAGGTCATGGAAGCTTCGTACGCT GCAGGTC 3'<br>**vip1_rev_int_devi**<br>TATTTAGTTTTGGGTTACTAAATTAAAAATTGGGTGTGATCACTAGCATAGGCCACTAGTG GATCTGA<br><br>**confirmation_vip1**<br>**5 prime confirmation**<br>**AGAAGAAGATTTACTCTCCCGTGAT**<br>**3 prime confirmation**<br>**CAAAGAATGGAAGAGAAATTTTGAA** | This Study |
| **Primer for deletion of *mot3***<br>**mot3_for_int_devi**<br>CAACAGTAGGCAAATAGTAAAGGGACATATCATATTTGAGCAATGGAAGCTTCGTACGCT GCAGGTC<br>**mot3_rev_int_devi**<br>AAATGAGTGGGAAGGGATATTTTGTGTGTCTATAAAGTCTATCTAGCATAGGCCACTAGT GGATCTGA<br>**Primer for confirmation of *mot3* deletion**<br>**1. 5 prime confirmation**<br>CTCCGTCTGGATTTACTAAACTTTG<br>GCCTCGAAACGTGAGTCTTTTC<br>**2. 3 prime confirmation**<br>AGTTTTCTCCTTCATTACAGAAAC<br>TGAATTCATCAAGAGATTTGAAACA | This Study |
| **Primer for deletion of *lrg1***<br>**lrg1_for_int_devi**<br>AGAGCAGACAAATTATCAAACAACAAGTACCGGAGGTGAGCAATGGAAGCTTCGTACGC TGCAGGTC<br>**lrg1_rev_int_devi**<br>AGAAAAAAAAGGAAAATGAGGGGAAACTTACAGTTTCTGCCTATTAGCATAGGCCACTAG TGGATCTGA<br>**Primer for confirmation of *lrg1* deletion**<br>**1. 5 prime confirmation for**<br>TCCCCACAAAGTATTTACTTCAAGA<br>GCCTCGAAACGTGAGTCTTTTC<br>**2. 3 prime confirmation rev**<br>AGTTTTCTCCTTCATTACAGAAAC | This Study |

| | |
|---|---|
| TTCTTCTGAAAGATTTCAATTTGCT | |
| **Primer for deletion of *mrn1***<br>**mrn1_for_int_devi**<br>TTTTTTCTTCACCATCACATACTACTTCAATTGCATTAAAACATGGAAGCTTCGTACGCTGCAGGTC<br>**mrn1_rev_int_devi**<br>ACTAAACATCTACGTACATACATATACATATATACATAATGTTTAGCATAGGCCACTAGTGGATCTGA<br>**Primer for confirmation of *lrg1* deletion**<br>**1. 5 prime confirmation**<br>ATTAGGCTCCTTAATTGGTTCAACT<br>GCCTCGAAACGTGAGTCTTTTC<br>**2. 3 prime confirmation**<br>AGTTTTCTCCTTCATTACAGAAAC<br>GTAAAATATTACCGCTTCGGAGATT | This Study |
| **seg2_for_int_devi**<br>AGGCAAGTATCAACAAATAGTGGGAGCATTGGAAATAGCGGGATGGAAGCTTCGTACGCTGCAGGTC<br>**seg2_rev_int_devi**<br>CTTACATTTCATTTTAGGTTTCCTATGTTTTCTTCTTTCAAATCAGCATAGGCCACTAGTGGATCTGA<br>**Primer for confirmation of *seg2* deletion**<br>**1. 5 prime confirmation for**<br>AATTCGAGAGAAAATAAATGGGAAG<br>**2. 3 prime confirmation rev**<br>CACCTTACAGTTCACCTTTGTCTTT | This Study |
| **Swa2_for_int_devi**<br>GCTTCTGGAAAGGACGCAGCCTGCAAGAAACAGTCAACATCAATGCGGGTCACCCGGCCAGCGACATGGAG<br>**Swa2_rev_int_devi**<br>ACATATCAAAAACAACTGAGCGAAGCAGGCACACAAGGGAAATCACGAATCGACAGCAGTATAGCGACCAGC<br>**Primer for confirmation of *seg2* deletion**<br>**1. 5 prime confirmation for**<br>AACAAGTGCAGGCTAACATACTTTC<br>**2. 3 prime confirmation rev**<br>ATTGATAATAATGCGCCTACAAAAA | This Study |
| **Ire1_Kan_For**<br>GAAAAATGCGTCTACTTCGAAGAAACATGTTAGTATTGACACTGCTCGTTTGGACATGGAGGCCCAGAATACCCTCC<br>**Ire1_Kan_Rev**<br>ATTAATGCAATAATCAACCAAGAAGAAGCAGAGGGGCATGAACATGATCGACAGCAGTATAGCGACCAGC<br>**Primer for confirmation of *ire1* deletion**<br>**1. 5 prime confirmation for**<br>AATAGGTTTTCGCTATTTTATTGCC<br>**2. 3 prime confirmation rev**<br>**TCACAAAGATTAAAGGAGCTATTGG** | This Study |
| **Kex2_for_int_devi**<br>TTGGCCTCGTCACATAATTATAAACTACTAACCCATTATCAGATGCGGGTCACCCGGCCAGCGACATGGAG<br>**Kex2_rev_int_devi** | This Study |

| | |
|---|---|
| AAAATGCTATTTTGTAATTTGAAGCTTTCTGTACATATCGAATCACGAATCGACAGCAGTA<br>TAGCGACCAGC<br>**Primer for confirmation of *kex2* deletion**<br>**1. 5 prime confirmation for**<br>ATAATCAATGAGGGTCATTTTCTGA<br>**2. 3 prime confirmation rev**<br>CGTTTTAAAGTTATTCAGCTGTGGT | |
| **Kex2_for_int_devi**<br>TTGGCCTCGTCACATAATTATAAACTACTAACCCATTATCAGATGCGGGTCACCCGGCCAG<br>CGACATGGAG<br>**Kex2_rev_int_devi**<br>AAAATGCTATTTTGTAATTTGAAGCTTTCTGTACATATCGAATCACGAATCGACAGCAGTA<br>TAGCGACCAGC<br>**Primer for confirmation of *kex2* deletion**<br>**1. 5 prime confirmation for**<br>ATAATCAATGAGGGTCATTTTCTGA<br>**2. 3 prime confirmation rev**<br>CGTTTTAAAGTTATTCAGCTGTGGT | This Study |
| **For Kanmx integration confirmation in selected deletions**<br>**conf_rev_5p_Kanmx_devi** GCCTCGAAACGTGAGTCTTTTC<br>**conf_for_3p_Kanmx_devi** AGTTTTCTCCTTCATTACAGAAAC | This Study |
| **ecm1_for_int_devi**<br>AGCTGAAAATTTTTTTGGTTAAGGACCCTTTAGAAGTATTGAATGGATGTCCACG<br>AGCTCTCTGAAGCTTCGTACGCTGCAGGTC<br>**ecm1_rev_int_devi**<br>ACAAGAATCGTGTATTCCTAGGACCTTTCATATGAAAATTTTCTAATCGATGAAT<br>TCGAGCTCGGCATAGGCCACTAGTGGATCTGA<br>**Primer for confirmation of *ecm1* deletion**<br>**1. 5 prime confirmation for**<br>ACGCCTAACAATCACGGTGAAAT<br>**2. 3 prime confirmation rev**<br>TCACACGGTTTTCAGCTAACCCATAA | This Study |

#### III. Kits and Enzymes Used

| REAGENT | SOURCE | Catalogue number |
|---|---|---|
| QIAamp DNA Mini Kit (250) | Qiagen | 51306 |
| Pierce™ BCA Protein Assay Kit | Thermo Fisher Scientific | 23225 |
| HiYield™ Gel/PCR DNA Mini Kit | RBC Real Biotech Corporation | QDF300 |
| Taq DNA Polymerase, recombinant | Thermo Scientific | |
| Deoxynucleotide Solution Mix (dNTPs) | Thermo Scientific | R0192 |
| Q5® High-Fidelity DNA Polymerase | New England Biolabs® (NEB®) | NEB #M0491 |
| Herculase II Fusion DNA Polymerases | Agilent Technologies | |

| | | |
|---|---|---|
| 100 bp DNA ladder | New England Biolabs® (NEB®) | N3231L |
| 1kb DNA ladder | New England Biolabs® (NEB®) | N3232S |

#### IV. Recombinant DNA

| REAGENT | SOURCE | IDENTIFIER |
|---|---|---|
| WTNAT in pRS316 | Wt TS5 in pRS316 fused with DsRed at C terminal construct along with URA were taken for making integrated TT strain and UE parental strain By4741 and yMJ003. | This study |
| TS22 in pRS316 | TS22 is a NAT variant having mutation E49G, V52M and D110V fused with DsRed at C terminal construct along with URA were taken for making integrated TT strains, UE parental strain By4741 and yMJ003. | This study |
| pFA6-kanMX4 | Used for gene deletion in yeast | This study |
| pUG6-loxP-KanMX-loxp | Used for gene deletion in yeast | This study |

#### v. Bacterial and Yeast Strains

| REAGENT | IDENTIFIER | SOURCE |
|---|---|---|
| **Bacterial Strains (*E. coli*)** | | |
| *E. coli* DH5α | F– endA1 glnV44 thi-1 recA1 relA1 gyrA96 deoR nupG purB20 φ80dlacZΔM15 Δ(lacZYA-argF)U169, hsdR17(rK–mK+), λ– | |
| **Yeast Strains (*S. cerevisiae*)** | | |
| BY4741 | (MAT*a* his3Δ1 leu2Δ0 lys2Δ0 ura3Δ0) | American Type culture collection (ATCC) |
| S288C | *MATα SUC2 gal2 mal2 mel flo1 flo8-1 hap1 ho bio1 bio6* | American Type culture collection (ATCC) |
| TT strains (49) | From BY4741(30) and S288C(19) background stains | This Study |
| Deletion strains used | ire1*Δ*, *inh1Δ, vip1Δ, seg2Δ, kex2Δ, mot3Δ, swa2Δ,mrn1Δ,lrg1Δ, ecm1Δ*<br>1. *BY4741* ire1*Δ::KanMX*<br>2. *BY4741 inh1Δ::KanMX*<br>3. *BY4741 vip1Δ::KanMX*<br>4. *BY4741 seg2Δ::KanMX*<br>5. *BY4741 kex2Δ::KanMX*<br>6. *BY4741 mot3Δ::KanMX*<br>7. *BY4741 mrn1Δ::KanMX* | Invitrogen |

| | | |
|---|---|---|
| | 8. *BY4741 lrg1Δ::KanMX* | |
| GFP integrated strains used | TDH2-GFP | Invitrogen |
| ire1 deletion in background of TT strains | TTB1, TTB2,TTB3,TTB5 (*ire1* replaced with *KanMX*) | This Study |
| Reconstructed deletions in the background of BY4741 (*inh1Δ, vip1Δ, seg2Δ, kex2Δ, mot3Δ, swa2Δ,mrn1Δ,lrg1Δ, ecm1Δ)* | 1. *BY4741 inh1Δ::KanMX*<br>2. *BY4741 vip1Δ::KanMX*<br>3. *BY4741 seg2Δ::KanMX*<br>4. *BY4741 kex2Δ::KanMX*<br>5. *BY4741 mot3Δ::KanMX*<br>6. *BY4741 mrn1Δ::KanMX*<br>7. *BY4741 lrg1Δ::KanMX*<br>8. *BY4741 ecm1Δ::KanMX* | This Study |

#### VI. Software and Algorithms

| | |
|---|---|
| Trimmomatic | **(Bolger et al., 2014)** |
| BWA | **(Li and Durbin, 2010)** |
| SAMtools | **(Li et al., 2009)** |
| GATK | **(McKenna et al., 2010)** |
| hisat2 | (**Kim et al., 2019)** |
| imageJ | **(Schneider et al., 2012)** |

All data analysis and plots were created using R and Python.

### References

Bolger AM, Lohse M, Usadel B. 2014. Trimmomatic: a flexible trimmer for Illumina sequence data. *Bioinformatics* **30**:2114–2120.

Huxley C, Green ED, Dunham I. 1990. Rapid assessment of S. cerevisiae mating type by PCR. *Trends* Genet 6:236.

Kahm M, Hasenbrink G, Lichtenberg-Fraté H, Ludwig J, Kschischo M. 2010. grofit: Fitting biological growth curves with R. Journal of Statistical Software 33:1–21.

Kim D, Paggi JM, Park C, Bennett C, Salzberg SL. 2019. Graph-based genome alignment and genotyping with HISAT2 and HISAT-genotype. *Nat Biotechnol* **37**:907–915.

Li H, Durbin R. 2010. Fast and accurate long-read alignment with Burrows–Wheeler transform. *Bioinformatics* **26**:589–595.

Li H, Handsaker B, Wysoker A, Fennell T, Ruan J, Homer N, Marth G, Abecasis G, Durbin R, Subgroup 1000 Genome Project Data Processing. 2009. The sequence alignment/map format and SAMtools. *bioinformatics* **25**:2078–2079.

McKenna A, Hanna M, Banks E, Sivachenko A, Cibulskis K, Kernytsky A, Garimella K, Altshuler D, Gabriel S, Daly M. 2010. The Genome Analysis Toolkit: a MapReduce framework for analyzing next-generation DNA sequencing data. *Genome Res* **20**:1297–1303.

### Supplemental Tables

**Table S1**: Details of different Thermotolerant Strains and their origin from parental background.

| Serial number | Well-positioned in the deep multiwell plate (well-named according to the combination of letters and numbers) | TT Strain name | Parental background<br><br>(Source of origin of TT strains) |
|---|---|---|---|
| 1 | A1 | TTA1 | S288C background |
| 2 | A2 | TTA2 | |
| 3 | A3 | TTA3 | |
| 4 | A4 | TTA4 | |
| 5 | A5 | TTA5 | |
| 6 | A6 | TTA6 | |
| 7 | A7 | TTA7 | |
| The wells such as A8, A9, A10, A11 and A12 Strains from the S288C background were lost during the ALE process. | | | |
| 8 | B1 | TTB1 | BY4741 background |
| 9 | B2 | TTB2 | |
| 10 | B3 | TTB3 | |
| 11 | B4 | TTB4 | |
| 12 | B5 | TTB5 | |
| 13 | B6 | TTB6 | |
| 14 | B7 | TTB7 | |
| 15 | B8 | TTB8 | |
| 16 | B9 | TTB9 | |
| 17 | B10 | TTB10 | |
| 18 | B11 | TTB11 | |
| In wells such as B12, Strains from the BY4741 background got cross-contaminated with S288C during the process of ALE. | | | |
| 19 | D1 | TTD1 | S288C background |
| 20 | D2 | TTD2 | |
| 21 | D3 | TTD3 | |

| | | | |
|---|---|---|---|
| 22 | D4 | TTD4 | |
| 23 | D5 | TTD5 | |
| 24 | D7 | TTD7 | |
| 25 | D8 | TTD8 | |
| In wells such as D6 and D10 Strains from the S288C background got cross-contaminated with BY4741 and in wells such as D9, D11, and D12 Strains from the S288C background were lost during the ALE process. | | | |
| 26 | E1 | TTE1 | BY4741 background |
| 27 | E2 | TTE2 | |
| 28 | E3 | TTE3 | |
| 29 | E4 | TTE4 | |
| 30 | E5 | TTE5 | |
| 31 | E6 | TTE6 | |
| 32 | E7 | TTE7 | |
| 33 | E8 | TTE8 | |
| 34 | E9 | TTE9 | |
| 35 | E10 | TTE10 | |
| In wells such as B11 Strains from the BY4741 background got cross-contaminated with S288C and in wells such as B12 Strains from the BY4741 background were lost during the ALE process. | | | |
| 37 | F3 | TTF3 | S288C background |
| 38 | F4 | TTF4 | |
| 39 | F5 | TTF5 | |
| The wells such as F1, F2 and F6 Strains from the S288C background were lost during the ALE process. | | | |
| 40 | F7 | TTF7 | BY4741 background |
| 41 | F8 | TTF8 | |
| 42 | F9 | TTF9 | |
| 43 | F12 | TTF12 | |
| The wells such as F10 and F11 Strains from the BY4741 background were lost during the ALE process. | | | |
| 44 | G2 | TTG2 | S288C background |
| 45 | G3 | TTG3 | |

| | | | |
|---|---|---|---|
| In wells such as G5 Strains from the S288C background got cross-contaminated with BY4741 and in wells such as G1,G4 and G6 Strains from the S288C background were lost during the ALE process. | | | |
| 46 | G9 | TTG9 | BY4741 background |
| 47 | G10 | TTG10 | |
| 48 | G11 | TTG11 | |
| 49 | G12 | TTG12 | |
| The wells such as G7 and G8 Strains from the BY4741 background were lost during the ALE process. | | | |
| The wells from C1-C12 and H1-H12 were taken as bank | | | |
| **Details of strain arrangements were given in the supplementary figure S1A.** | | | |

**Supplement TableS2:** List of missense/nonsense mutations in TT strains in BY4741 and S288C parental background strains

| **List of missense mutations in TT strains in BY4741** | | | | |
|---|---|---|---|---|
| Chr_position | Annotation | Mutation | AA change | Thermotolerant strains having the mutations |
| chr01_377 | YAL069W | c.43A>C | p.Thr15Pro | TTB6 TTB11 |
| chr01_402 | YAL069W | c.68C>T | p.Pro23Leu | TTB11 |
| chr01_509 | YAL069W | c.175G>A | p.Glu59Lys | TTB2 TTE5 |
| chr01_528 | YAL069W | c.194T>C | p.Leu65Pro | TTB2 |
| chr01_540 | YAL068W-A | c.3G>T | p.Met1? | TTB5 |
| chr01_568 | YAL068W-A | c.31C>T | p.Pro11Ser | TTE5 |
| chr01_602 | YAL069W | c.268C>T | p.His90Tyr | TTE10 TTE10_C1 |
| chr01_610 | YAL068W-A | c.73G>A | p.Asp25Asn | TTE10 TTE10_C1 |
| chr01_627 | YAL069W | c.293C>T | p.Ser98Leu | TTE10 TTE10_C1 |
| chr01_633 | YAL069W | c.299T>C | p.Val100Ala | TTE10 TTE10_C1 |
| chr01_636 | YAL069W | c.302C>A | p.Pro101Gln | TTE10 TTE10_C1 |
| chr01_25955 | YAL063C | c.2014A>G | p.Ile672Val | TTB4 TTB7 TTB8 TTE1 TTE3 TTE4 TTE6 TTE7 TTE8 TTE7_C1 TTG10_C1 TTG9 TTG10 TTG12 |
| chr01_26839 | YAL063C | c.1130T>C | p.Phe377Ser | TTB4 TTE5 TTF12 TTG10_C1 TTG10 TTG11 TTG12 |
| chr01_26846 | YAL063C | c.1123A>G | p.Ser375Gly | TTE5 TTF12 0 TTG10 TTG11 TTG12 |
| chr01_26848 | YAL063C | c.1121A>C | p.Asn374Thr | TTE5 TTF12 TTG10_C1 TTG10 TTG11 TTG12 |
| chr01_62514 | YAL042W | c.1199A>C | p.Lys400Thr | TTB1 |
| chr01_204390 | YAR050W | c.986A>C | p.Asn329Thr | TTB9 |
| chr01_204392 | YAR050W | c.988A>G | p.Ser330Gly | TTB9 |
| chr01_204419 | YAR050W | c.1015T>A | p.Leu339Met | TTB9 |
| chr01_204428 | YAR050W | c.1024G>A | p.Val342Ile | TTB9 |
| chr01_204442 | YAR061W | c.64A>G | p.Ile22Val | TTB9 |
| chr01_204450 | YAR050W | c.1046G>C | p.Arg349Pro | TTB9 |
| chr01_204472 | YAR061W | c.34T>A | p.Tyr12Asn | TTB9 |
| chr01_204473 | YAR050W | c.1069A>G | p.Ile357Val | TTB9 |
| chr01_204477 | YAR050W | c.1073G>A | p.Arg358Lys | TTB9 |
| chr01_206272 | YAR061W | c.13A>G | p.Thr5Ala | TTB9 |
| chr01_206293 | YAR061W | c.34T>C | p.Phe12Leu | TTB9 |
| chr01_206296 | YAR061W | c.37G>T | p.Asp13Tyr | TTB9 |
| chr01_206299 | YAR061W | c.40T>G | p.Phe14Val | TTB9 |
| chr01_206302 | YAR061W | c.43C>T | p.His15Tyr | TTB9 |
| chr01_206309 | YAR050W | c.2905G>A | p.Val969Ile | TTB9 |
| chr01_206311 | YAR061W | c.52T>G | p.Tyr18Asp | TTB9 |
| chr01_206314 | YAR061W | c.55C>T | p.His19Tyr | TTB9 |
| chr01_206317 | YAR061W | c.58C>T | p.His20Tyr | TTB9 |
| chr01_206326 | YAR061W | c.67A>C | p.Asn23His | TTB9 |
| chr01_206336 | YAR050W | c.2932C>G | p.Gln978Glu | TTB9 |
| chr01_206337 | YAR050W | c.2933A>T | p.Gln978Leu | TTB9 |
| chr02_408 | YBL113C | c.2251A>G | p.Asn751Asp | TTB9 |
| chr02_465 | YBL113C | c.2194A>G | p.Met732Val | TTB11 |
| chr02_1187 | YBL113C | c.1472G>A | p.Gly491Glu | TTB4 TTB11 TTE3 TTE4 TTF7 TTG12 |

| | | | | |
|---|---|---|---|---|
| chr02_1254 | YBL113C | c.1405A>G | p.Ile469Val | TTB4 TTE3 TTE4 TTF7 TTG11 TTG12 |
| chr02_1372 | YBL113C | c.1287C>G | p.Phe429Leu | TTB4 TTB7 TTE4 TTG12 |
| chr02_1596 | YBL113C | c.1063G>A | p.Asp355Asn | TTG12 |
| chr02_1619 | YBL113C | c.1040G>A | p.Ser347Asn | TTG12 |
| chr02_1625 | YBL113C | c.1034A>G | p.Asn345Ser | TTG12 |
| chr02_1691 | YBL113C | c.968A>G | p.Asn323Ser | TTB7 TTB11 TTG11 |
| chr02_1778 | YBL113C | c.881T>C | p.Ile294Thr | TTB7 TTF7 TTG12 |
| chr02_1785 | YBL113C | c.874G>A | p.Glu292Lys | TTB11 |
| chr02_37269 | YBL100C | c.35C>T | p.Ser12Leu | TTB6 |
| chr02_37891 | YBL099W | c.839A>T | p.Gln280Leu | TTB1 |
| chr02_38066 | YBL099W | c.1014G>T | p.Leu338Phe | TTB11 |
| chr02_38262 | YBL099W | c.1210G>T | p.Val404Phe | TTB10 |
| chr02_316450 | YBR039W | c.876A>T | p.Arg292Ser | TTB5 |
| chr02_316451 | YBR039W | c.877C>A | p.Gln293Lys | TTE9 |
| chr02_316454 | YBR039W | c.880G>C | p.Ala294Pro | TTB4 |
| chr02_316493 | YBR039W | c.919G>T | p.Ala307Ser | TTE1 |
| chr02_511204 | YBR136W | c.5536G>A | p.Glu1846Lys | TTE1 |
| chr02_676843 | YBR229C | c.2380G>T | p.Ala794Ser | TTE10 TTE10_C1 |
| chr02_781030 | YBR289W | c.1363G>A | p.Glu455Lys | TTE9 |
| chr02_797431 | YBR296C | c.1093G>T | p.Gly365Cys | TTF7 |
| chr03_188372 | YCR038C | c.599G>A | p.Arg200Lys | TTB7 |
| chr03_226636 | YCR068W | c.107C>G | p.Pro36Arg | TTB7 |
| chr04_1875 | YDL248W | c.74T>C | p.Ile25Thr | TTE9 TTG10 |
| chr04_2178 | YDL248W | c.377T>C | p.Val126Ala | TTG11 TTG12 |
| chr04_135222 | YDL181W | c.44G>A | p.Arg15His | TTB5 |
| chr04_451037 | YDR001C | c.1439C>A | p.Thr480Lys | TTE9 |
| chr04_580147 | YDR064W | c.151G>A | p.Gly51Ser | TTE7_C1 |
| chr04_610774 | YDR082W | c.334G>T | p.Asp112Tyr | TTB5 |
| chr04_973235 | YDR258C | c.1009C>T | p.Arg337Cys | TTB7 |
| chr04_1220727 | YDR371W | c.1315A>T | p.Asn439Tyr | TTB5 |
| chr04_1226357 | YDR375C | c.180C>G | p.Ile60Met | TTE7_C1 |
| chr04_1395353 | YDR466W | c.233T>A | p.Leu78Gln | TTB1 |
| chr04_1525404 | YDR544C | c.119C>A | p.Pro40His | TTB8 TTB11 TTG10_C1 TTG10 |
| chr04_1525450 | YDR544C | c.73C>A | p.Pro25Thr | TTB6 |
| chr04_1525464 | YDR544C | c.59A>C | p.His20Pro | TTB1 TTB2 TTB3 TTB7 TTB8 TTB9 TTE2 TTE3 TTE4 TTE10 TTF7 TTF8 TTE10_C1 TTE7_C1 TTG12 |
| chr05_385 | YEL077C | c.3713A>G | p.Lys1238Arg | TTB1 TTE1 TTE6 TTB10 |
| chr05_998 | YEL077C | c.3100A>G | p.Arg1034Gly | TTB7 TTE10_C1 |
| chr05_1238 | YEL077C | c.2860A>G | p.Ile954Val | TTE10_C1 |
| chr05_1544 | YEL077C | c.2554A>G | p.Asn852Asp | TTE6 |
| chr05_2084 | YEL077C | c.2014A>G | p.Asn672Asp | TTE1 TTG12 |
| chr05_233432 | YER041W | c.2069A>C | p.Glu690Ala | TTF8 |
| chr05_252314 | YER049W | c.1684G>T | p.Asp562Tyr | TTE3 |
| chr05_451297 | YER140W | c.830A>T | p.Lys277Ile | TTB11 |
| chr05_532101 | YER172C | c.2824A>T | p.Thr942Ser | TTG12 |
| chr05_542085 | YER176W | c.1493A>T | p.Asn498Ile | TTE7_C1 |
| chr05_544565 | YER177W | c.52G>A | p.Ala18Thr | TTE10_C1 |
| chr05_570283 | YER189W | c.226A>G | p.Ile76Val | TTB9 |

| | | | | |
|---|---|---|---|---|
| chr06_986 | YFL067W | c.151C>A | p.Arg51Ser | TTB1 TTB5 TTB7 TTE1 TTE6 TTE10 TTE10_C1 TTE7_C1 |
| chr06_1022 | YFL067W | c.187A>C | p.Ser63Arg | TTB5 TTE6 TTF7 TTF9 TTG10 TTG11 |
| chr06_5068 | YFL063W | c.3G>A | p.Met1? | TTB2 TTE6 |
| chr06_5135 | YFL063W | c.70A>T | p.Ile24Phe | TTE9 |
| chr06_5153 | YFL063W | c.88C>G | p.Arg30Gly | TTE9 |
| chr06_5410 | YFL063W | c.345T>A | p.Phe115Leu | TTB2 |
| chr06_6847 | YFL062W | c.422T>C | p.Ile141Thr | TTE6 TTE7 |
| chr06_41819 | YFL047W | c.1399C>T | p.Pro467Ser | TTF7 |
| chr06_128959 | YFL007W | c.5481A>C | p.Glu1827Asp | TTB11 |
| chr06_208777 | YFR028C | c.1292G>A | p.Gly431Asp | TTE8 |
| chr07_34099 | YGL249W | c.1002T>G | p.Asn334Lys | TTG11 |
| chr07_181558 | YGL172W | c.859C>A | p.Gln287Lys | TTE8 |
| chr07_292121 | YGL115W | c.89C>A | p.Thr30Lys | TTE10 TTE10_C1 |
| chr07_392689 | YGL059W | c.467G>A | p.Arg156Lys | TTB10 |
| chr07_437755 | YGL031C | c.180G>C | p.Lys60Asn | TTB1 |
| chr07_455805 | YGL021W | c.1021T>G | p.Ser341Ala | TTE7_C1 |
| chr07_481797 | YGL008C | c.870G>C | p.Leu290Phe | TTB7 |
| chr07_623634 | YGR067C | c.1153A>T | p.Asn385Tyr | TTB7 |
| chr07_677453 | YGR096W | c.833G>T | p.Gly278Val | TTE10_C1 |
| chr07_763153 | YGR137W | c.266G>A | p.Ser89Asn | TTB4 |
| chr07_992669 | YGR250C | c.853G>C | p.Val285Leu | TTE7_C1 |
| chr08_1569 | YHL050C | c.970C>T | p.Pro324Ser | TTB9 TTE4 TTF8 TTF12 |
| chr08_1712 | YHL050C | c.827A>G | p.Asn276Ser | TTG10_C1 TTG10 |
| chr08_1717 | YHL050C | c.822C>G | p.Ser274Arg | TTG10_C1 TTG10 |
| chr08_1748 | YHL050C | c.791A>G | p.Asn264Ser | TTF8 |
| chr08_1761 | YHL050C | c.778A>G | p.Asn260Asp | TTF8 |
| chr08_1835 | YHL050C | c.704T>C | p.Ile235Thr | TTG10 |
| chr08_1910 | YHL048W | MODIFIER | c.-4491G>A | TTB9 |
| chr08_1965 | YHL048W | MODIFIER | c.-4436C>T | TTB9 TTF12 |
| chr08_2068 | YHL048W | MODIFIER | c.-4333A>T | TTB4 TTB9 TTF12 |
| chr08_13664 | YHL044W | c.100C>G | p.Pro34Ala | TTE1 |
| chr08_164274 | YHR027C | c.436T>A | p.Phe146Ile | TTE8 |
| chr08_244131 | YHR073W | c.1552G>A | p.Ala518Thr | TTE7_C1 |
| chr08_367801 | YHR131C | c.90G>A | p.Met30Ile | TTB1 |
| chr08_426644 | YHR164C | c.2533A>C | p.Thr845Pro | TTG12 |
| chr08_556659 | YHR217C | c.382A>G | p.Thr128Ala | TTG12 |
| chr08_556668 | YHR217C | c.373A>T | p.Asn125Tyr | TTG12 |
| chr08_556833 | YHR217C | c.208A>C | p.Thr70Pro | TTB9 |
| chr08_556953 | YHR217C | c.88C>A | p.Pro30Thr | TTB2 TTB3 TTB7 TTB11 TTE4 TTE5 TTE6 TTE7 TTE9 TTF7 TTF8 TTF9 TTF12 TTB10 TTG10 TTG11 TTG12 |
| chr08_556965 | YHR217C | c.76C>A | p.Pro26Thr | TTB11 TTE7 TTG10 |
| chr08_560556 | YHR219W | c.386A>G | p.Asn129Ser | TTB11 TTG12 |
| chr08_560562 | YHR219W | c.392A>G | p.Asn131Ser | TTF8 |
| chr08_560729 | YHR219W | c.559G>A | p.Asp187Asn | TTB10 |
| chr08_560742 | YHR219W | c.572G>A | p.Ser191Asn | TTE10 TTE10_C1 TTG12 |
| chr09_68782 | YIL149C | MODIFIER | c.-714A>C | TTE10 |
| chr09_89565 | YIL138C | c.152A>T | p.Glu51Val | TTE10 TTE10_C1 |
| chr09_112024 | YIL129C | c.1218T>G | p.Asn406Lys | TTB6 |
| chr09_125528 | YIL125W | c.2836G>T | p.Val946Phe | TTB2 |

| | | | | |
|---|---|---|---|---|
| chr09_150659 | YIL113W | c.97A>G | p.Thr33Ala | TTB10 |
| chr10_93447 | YJL176C | c.1084T>A | p.Phe362Ile | TTB9 |
| chr10_94550 | YJL175W | c.502A>T | p.Ile168Phe | TTE3 |
| chr10_95673 | YJL174W | c.582G>T | p.Met194Ile | TTE1 |
| chr10_162283 | YJL132W | c.370G>T | p.Asp124Tyr | TTG9 |
| chr10_344754 | YJL050W | c.2233G>C | p.Gly745Arg | TTE7_C1 |
| chr10_368001 | YJL041W | c.2100C>G | p.Ile700Met | TTE5 |
| chr10_648235 | YJR121W | c.629C>T | p.Ala210Val | TTE3 |
| chr10_648378 | YJR121W | c.772C>T | p.Pro258Ser | TTB9 |
| chr10_648645 | YJR121W | c.1039G>T | p.Ala347Ser | TTE5 |
| chr10_715068 | YJR151C | c.673C>T | p.Pro225Ser | TTG10 |
| chr10_715122 | YJR151C | c.619C>T | p.Pro207Ser | TTG10_C1 TTG9 |
| chr10_720819 | YJR152W | c.1154T>A | p.Phe385Tyr | TTE10_C1 |
| chr10_738075 | YJR160C | c.1742C>A | p.Thr581Asn | TTB4 TTF7 TTG12 |
| chr10_745053 | YJR162C | c.209C>A | p.Thr70Asn | TTF7 |
| chr11_673 | YKL225W | c.223A>G | p.Ile75Val | TTG9 |
| chr11_689 | YKL225W | c.239T>A | p.Leu80His | TTG9 |
| chr11_704 | YKL225W | c.254T>C | p.Ile85Thr | TTG9 |
| chr11_778 | YKL225W | c.328C>T | p.Leu110Phe | TTF7 |
| chr11_51771 | YKL205W | c.1721A>T | p.Asp574Val | TTF12 |
| chr11_195466 | YKL130C | c.566G>T | p.Gly189Val | TTE3 |
| chr11_275437 | YKL088W | c.152G>T | p.Arg51Ile | TTB6 |
| chr11_374758 | YKL033W-A | c.251T>C | p.Leu84Pro | TTE1 |
| chr11_383842 | YKL029C | c.887G>A | p.Gly296Asp | TTE3 |
| chr11_495101 | YKR028W | c.842T>A | p.Ile281Lys | TTE7 |
| chr11_545049 | YKR054C | c.2880G>T | p.Trp960Cys | TTB11 |
| chr11_634818 | YKR098C | c.365C>A | p.Ala122Glu | TTF8 |
| chr12_5683 | YLL066W-B | c.79C>A | p.Pro27Thr | TTF9 |
| chr12_55850 | YLL040C | c.7796G>A | p.Gly2599Asp | TTB1 |
| chr12_141928 | YLL004W | c.856G>A | p.Asp286Asn | TTG9 |
| chr12_284539 | YLR077W | c.669G>T | p.Leu223Phe | TTE7_C1 |
| chr12_295403 | YLR083C | c.691C>A | p.Arg231Ser | TTF8 |
| chr12_317080 | YLR088W | c.975G>T | p.Met325Ile | TTB5 |
| chr12_490442 | YLR162W-A | c.38A>G | p.Asp13Gly | TTE5 |
| chr12_490486 | YLR162W-A | c.82T>C | p.Ser28Pro | TTE5 |
| chr12_522959 | YLR183C | MODIFIER | c.-948A>C | TTB2 TTB3 |
| chr12_556962 | YLR207W | c.176T>G | p.Phe59Cys | TTG12 |
| chr12_692061 | YLR274W | c.508T>C | p.Ser170Pro | TTB8 |
| chr12_799282 | YLR336C | c.716A>G | p.Asp239Gly | TTB6 |
| chr12_895044 | YLR387C | c.230T>C | p.Met77Thr | TTG11 |
| chr12_937951 | YLR410W | c.2810T>C | p.Val937Ala | TTE8 |
| chr12_1007819 | YLR436C | c.1028C>T | p.Ser343Phe | TTB5 |
| chr12_1069214 | YLR466C-B | c.98T>C | p.Ile33Thr | TTE10 TTE10_C1 TTG12 |
| chr12_1069244 | YLR466C-B | c.68C>T | p.Ser23Phe | TTE10 |
| chr12_1069251 | YLR466C-B | c.61A>C | p.Thr21Pro | TTE10 |
| chr12_1069253 | YLR466C-B | c.59C>A | p.Pro20His | TTE10 |
| chr13_11130 | YML131W | c.933T>A | p.Asp311Glu | TTE7_C1 |
| chr13_269181 | YMR001C | c.1953A>C | p.Lys651Asn | TTB11 |
| chr13_274156 | YMR004W | c.143C>T | p.Thr48Ile | TTG12 |
| chr13_676113 | YMR206W | c.221A>G | p.Glu74Gly | TTB6 |
| chr14_6669 | YNL338W | c.109C>A | p.Pro37Thr | TTE10 TTE10_C1 |
| chr14_6705 | YNL338W | c.145C>A | p.Pro49Thr | TTB8 |

| | | | | |
|---|---|---|---|---|
| chr14_16619 | YNL331C | c.630G>T | p.Trp210Cys | TTB5 |
| chr14_93244 | YNL287W | c.1251C>A | p.Asn417Lys | TTE5 |
| chr14_96315 | YNL285W | c.143G>T | p.Cys48Phe | TTE3 |
| chr14_203017 | YNL238W | c.590T>C | p.Phe197Ser | TTB2 TTB3 |
| chr14_203445 | YNL238W | c.1018G>T | p.Asp340Tyr | TTF8 |
| chr14_203477 | YNL238W | c.1050A>C | p.Glu350Asp | TTE3 |
| chr14_203949 | YNL238W | c.1522G>T | p.Val508Phe | TTB6 |
| chr14_204105 | YNL238W | c.1678T>C | p.Trp560Arg | TTB8 |
| chr14_364333 | YNL139C | c.1382A>C | p.Glu461Ala | TTF8 |
| chr14_468780 | YNL085W | c.1652T>C | p.Phe551Ser | TTB6 |
| chr14_475557 | YNL082W | c.2169G>T | p.Gln723His | TTB6 |
| chr14_575339 | YNL032W | c.836T>C | p.Leu279Pro | TTE5 |
| chr14_671253 | YNR023W | c.838G>T | p.Gly280Cys | TTB8 TTG9 TTG11 TTG12 |
| chr14_729814 | YNR055C | c.371C>T | p.Ala124Val | TTF9 |
| chr14_737251 | YNR059W | c.451T>G | p.Phe151Val | TTG9 |
| chr14_747114 | YNR063W | c.174A>T | p.Lys58Asn | TTF9 |
| chr14_770924 | YNR071C | c.543G>C | p.Leu181Phe | TTB8 |
| chr15_1832 | YOL165C | c.247G>T | p.Val83Phe | TTB7 |
| chr15_271061 | YOL028C | c.310G>A | p.Gly104Ser | TTB5 |
| chr15_460945 | YOR071C | c.332G>A | p.Arg111Lys | TTF7 |
| chr15_643443 | YOR164C | c.890C>A | p.Thr297Lys | TTE1 |
| chr15_856453 | YOR290C | c.3616G>T | p.Val1206Phe | TTE4 |
| chr15_857805 | YOR290C | c.2264T>A | p.Ile755Asn | TTB1 TTB11 TTE5 TTE6 TTE7 TTE7_C1 TTB10 |
| chr15_858615 | YOR290C | c.1454C>T | p.Pro485Leu | TTE1 |
| chr15_942127 | YOR330C | c.1069G>A | p.Glu357Lys | TTE2 |
| chr16_70920 | YPL254W | c.1436G>T | p.Trp479Leu | TTE10 TTE10_C1 |
| chr16_93960 | YPL242C | c.1150C>T | p.Arg384Cys | TTF8 |
| chr16_177346 | YPL195W | c.1124T>C | p.Val375Ala | TTE4 |
| chr16_191772 | YPL188W | c.367C>T | p.Pro123Ser | TTE9 |
| chr16_196234 | YPL184C | c.1555G>T | p.Gly519Cys | TTE10 TTE10_C1 |
| chr16_196680 | YPL184C | c.1109G>T | p.Arg370Ile | TTB9 |
| chr16_236855 | YPL167C | c.253A>G | p.Lys85Glu | TTF7 |
| chr16_270343 | YPL150W | c.2156C>A | p.Ala719Asp | TTE10_C1 |
| chr16_395408 | YPL084W | c.1371A>T | p.Glu457Asp | TTB2 |
| chr16_774844 | YPR120C | c.339A>C | p.Glu113Asp | TTB6 |
| chr16_943950 | YPR203W | c.71A>G | p.Lys24Arg | TTE7 |
| **(B): List of Nonsense mutations in TT strains in BY4741** | | | | |
| Chr_position | annotation | Mutation | AA change | TT strains |
| chr04_24338 | YDL240W | c.1516A>T | p.Arg506* | TTG10_C1 TTG10 |
| chr04_135192 | YDL181W | c.14C>A | p.Ser5* | TTE7 |
| chr04_1107311 | YDR320C | c.791C>A | p.Ser264* | TTG9 |
| chr04_1107362 | YDR320C | c.740C>A | p.Ser247* | TTE8 |
| chr04_1107741 | YDR320C | c.361G>T | p.Glu121* | TTE4 |
| chr04_1253344 | YDR389W | c.808C>T | p.Gln270* | TTF9 |
| chr11_239518 | YKL105C | c.3069C>A | p.Cys1023* | TTB6 |
| chr11_655183 | YKR103W | c.2101G>T | p.Gly701* | TTE2 |
| chr12_126866 | YLL012W | c.1333C>T | p.Gln445* | TTB5 |
| chr12_935592 | YLR410W | c.451G>T | p.Gly151* | TTB5 TTB6 TTB8 TTE4 TTE10 TTF8 TTF12 TTG9 TTF9 TTG11 TTG12 |
| chr13_257192 | YML006C | c.1222C>T | p.Gln408* | TTE8 |

| | | | | |
|---|---|---|---|---|
| chr13_363793 | YMR047C | c.2910C>A | p.Tyr970* | TTB7 |
| chr14_202853 | YNL238W | c.426G>A | p.Trp142* | TTG12 |
| chr14_203475 | YNL238W | c.1048G>T | p.Glu350* | TTB4 |
| chr15_291978 | YOL018C | c.97C>T | p.Gln33* | TTF7 |
| chr16_197624 | YPL184C | c.165T>A | p.Tyr55* | TTB11 TTE6 TTE7 |
| **(C): List of missense mutations in TT strains in S288C** | | | | |
| Chr_position | annotation | Mutation | AA change | TT strains |
| chr_I_377 | YAL069W | c.43A>C | p.Thr15Pro | TTG5 |
| chr_I_25955 | YAL063C | c.2014A>G | p.Ile672Val | TTG5 TTD5 TTD3 TTD2 TTD1 TTA7 |
| chr_I_26846 | YAL063C | c.1123A>G | p.Ser375Gly | TTA7 TTA5 TTA2 TTA2 |
| chr_I_26848 | YAL063C | c.1121A>C | p.Asn374Thr | TTA7 TTA5 TTA2 TTA2 |
| chr_I_103586 | YAL024C | c.2287C>A | p.His763Asn | TTA5 |
| chr_I_126383 | YAL016W | c.1505A>C | p.Lys502Thr | TTG5 |
| chr_I_206307 | YAR050W | c.2905G>A | p.Val969Ile | TTA7 |
| chr_II_408 | YBL113C | c.2251A>G | p.Asn751Asp | TTA7 |
| chr_II_796 | YBL113W-A | c.151T>C | p.Phe51Leu | TTD4 |
| chr_II_1187 | YBL113C | c.1472G>A | p.Gly491Glu | TTD4 TTA7 |
| chr_II_1254 | YBL113C | c.1405A>G | p.Ile469Val | TTD4 TTA7 |
| chr_II_1691 | YBL113C | c.968A>G | p.Asn323Ser | TTA7 |
| chr_II_38069 | YBL099W | c.1017T>G | p.His339Gln | TTG5 |
| chr_II_38262 | YBL099W | c.1210G>T | p.Val404Phe | TTD1 |
| chr_II_315922 | YBR039W | c.348G>T | p.Leu116Phe | TTF5 |
| chr_II_316449 | YBR039W | c.875G>A | p.Arg292Lys | TTA6 TTA5 TTA3 TTA3 TTA2 TTA1 TTA2 |
| chr_II_316451 | YBR039W | c.877C>A | p.Gln293Lys | TTF3 |
| chr_II_646439 | YBR211C | c.694G>T | p.Ala232Ser | TTF5 |
| chr_II_781281 | YBR289W | c.1615G>A | p.Gly539Ser | TTF4 |
| chr_II_795843 | YBR295W | c.2995A>C | p.Lys999Gln | TTF4 |
| chr_IV_135195 | YDL181W | c.17C>A | p.Ala6Glu | TTG2 |
| chr_IV_450521 | YDR001C | c.1955G>A | p.Arg652Lys | TTF5 |
| chr_IV_499069 | YDR028C | c.1811C>A | p.Thr604Asn | TTG2 |
| chr_IV_500198 | YDR028C | c.682C>A | p.Leu228Ile | TTD7 TTD5 |
| chr_IV_611662 | YDR082W | c.1222G>T | p.Ala408Ser | TTA5 |
| chr_IV_642743 | YDR097C | c.1095A>C | p.Gln365His | TTD7 TTD5 |
| chr_IV_948960 | YDR243C | c.1325T>A | p.Phe442Tyr | TTF4 |
| chr_IV_949247 | YDR243C | c.1038G>A | p.Met346Ile | TTF4 |
| chr_IV_1296209 | YDR414C | c.478A>T | p.Ile160Phe | TTA6 TTA5 TTA3 TTA3 TTA2 TTA1 TTA2 |
| chr_IV_1511040 | YDR537C | c.422C>T | p.Ala141Val | TTG5 |
| chr_IV_1525406 | YDR544C | c.118C>T | p.Pro40Ser | TTD5 |
| chr_IV_1525465 | YDR544C | c.59A>C | p.His20Pro | TTG2 TTD8 TTD5 TTD1 TTA7 TTA6 TTA5 TTA3 TTA3 TTA2 TTA1 TTA2 |
| chr_IX_78726 | YIL144W | c.653T>A | p.Phe218Tyr | TTD4 TTD3 TTD2 TTA7 |
| chr_IX_109321 | YIL129C | c.3917T>C | p.Phe1306Ser | TTD8 |
| chr_IX_112020 | YIL129C | c.1218T>G | p.Asn406Lys | TTG5 |
| chr_V_385 | YEL077C | c.3713A>G | p.Lys1238Arg | TTG3 TTA7 |
| chr_V_1238 | YEL077C | c.2860A>G | p.Ile954Val | TTG2 TTA7 |
| chr_V_1544 | YEL077C | c.2554A>G | p.Asn852Asp | TTA2 TTA1 TTA2 |
| chr_V_533192 | YER172C | c.2830C>G | p.Leu944Val | TTA7 |
| chr_VI_899 | YFL067W | c.64G>A | p.Ala22Thr | TTG5 |

| | | | | |
|---|---|---|---|---|
| chr_VI_5180 | YFL063W | c.115G>T | p.Val39Leu | TTA7 |
| chr_VI_178696 | YFR016C | c.2048G>A | p.Gly683Glu | TTF4 |
| chr_VI_214570 | YFR030W | c.1259C>T | p.Ala420Val | TTD2 |
| chr_VII_114369 | YGL203C | c.296C>T | p.Pro99Leu | TTD1 |
| chr_VIII_435539 | YHR165C | c.1410G>T | p.Leu470Phe | TTD1 |
| chr_VIII_499958 | YHR201C | c.1186G>A | p.Glu396Lys | TTG5 |
| chr_VIII_556955 | YHR217C | c.88C>A | p.Pro30Thr | TTD5 TTA5 TTA2 TTA2 |
| chr_VIII_556967 | YHR217C | c.76C>A | p.Pro26Thr | |
| chr_X_92838 | YJL176C | c.1693T>G | p.Phe565Val | TTD1 |
| chr_X_268809 | YJL088W | c.11C>T | p.Thr4Ile | TTF5 |
| chr_X_648481 | YJR121W | c.875T>A | p.Phe292Tyr | TTG2 |
| chr_X_648650 | YJR121W | c.1044T>A | p.Asp348Glu | TTG3 |
| chr_X_648651 | YJR121W | c.1045G>A | p.Asp349Asn | TTD8 TTD7 TTD5 |
| chr_XI_509 | YKL225W | c.59G>T | p.Ser20Ile | TTF3 |
| chr_XI_546 | YKL225W | c.96T>G | p.Ile32Met | TTF3 |
| chr_XI_499797 | YKR029C | c.37G>T | p.Asp13Tyr | TTF5 |
| chr_XII_5741 | YLL066W-B | c.137T>C | p.Phe46Ser | TTG5 TTG2 TTD5 TTD4 TTD1 TTA2 TTA1 TTA2 |
| chr_XII_5747 | YLL066W-B | c.143T>G | p.Leu48Arg | TTA7 |
| chr_XII_291386 | YLR081W | c.1175A>G | p.His392Arg | TTG5 |
| chr_XII_938945 | YLR410W | c.1405C>G | p.Gln469Glu | TTF5 |
| chr_XII_938958 | YLR410W | c.1418G>T | p.Gly473Val | TTF4 |
| chr_XIII_161077 | YML057W | c.898G>T | p.Asp300Tyr | TTF4 |
| chr_XIII_410255 | YMR070W | c.1102C>T | p.His368Tyr | TTA6 |
| chr_XIII_450454 | YMR091C | c.912T>G | p.Asn304Lys | TTA7 |
| chr_XIII_470447 | YMR102C | c.1906G>A | p.Glu636Lys | TTG2 |
| chr_XIII_499621 | YMR116C | c.795G>T | p.Leu265Phe | TTA2 TTA2 |
| chr_XIII_630505 | YMR185W | c.1481A>G | p.Asp494Gly | TTG5 |
| chr_XIII_836548 | YMR283C | c.320G>T | p.Arg107Ile | TTG3 |
| chr_XIII_871383 | YMR302C | c.1243A>G | p.Lys415Glu | TTD2 |
| chr_XIV_33830 | YNL322C | c.406G>A | p.Val136Ile | TTD8 |
| chr_XIV_136760 | YNL270C | c.902C>T | p.Pro301Leu | TTG5 |
| chr_XIV_202549 | YNL238W | c.122G>T | p.Ser41Ile | TTF4 |
| chr_XIV_203085 | YNL238W | c.658G>A | p.Glu220Lys | TTD4 TTD3 TTD2 TTA7 |
| chr_XIV_203215 | YNL238W | c.788T>A | p.Leu263Gln | TTD8 |
| chr_XIV_203258 | YNL238W | c.831T>G | p.Asp277Glu | TTA2 TTA1 TTA2 |
| chr_XIV_203367 | YNL238W | c.940A>C | p.Asn314His | TTF3 |
| chr_XIV_203400 | YNL238W | c.973G>A | p.Asp325Asn | TTA5 |
| chr_XIV_203650 | YNL238W | c.1223G>T | p.Arg408Ile | TTA6 TTA3 TTA3 |
| chr_XIV_203949 | YNL238W | c.1522G>T | p.Val508Phe | TTG5 |
| chr_XIV_334406 | YNL161W | c.1810T>G | p.Cys604Gly | TTA7 |
| chr_XIV_468782 | YNL085W | c.1652T>C | p.Phe551Ser | TTG5 |
| chr_XIV_475559 | YNL082W | c.2169G>T | p.Gln723His | TTG5 |
| chr_XV_245970 | YOL045W | c.2474C>T | p.Pro825Leu | TTA7 |
| chr_XV_265685 | YOL032W | c.257A>G | p.Gln86Arg | TTD3 |
| chr_XV_545930 | YOR117W | c.902T>A | p.Val301Glu | TTD8 |
| chr_XV_557930 | YOR124C | c.713C>A | p.Thr238Asn | TTG5 |
| chr_XV_773264 | YOR231W | c.664G>C | p.Glu222Gln | TTG5 |
| chr_XVI_101163 | YPL237W | c.668A>G | p.Glu223Gly | TTG5 |
| chr_XVI_893401 | YPR178W | c.1070G>T | p.Trp357Leu | TTF5 |

| (D): List of Nonsense mutations in TT strains in S288C | | | | |
|---|---|---|---|---|
| Chr_position | annotation | Mutation | AA change | TT strains |
| chr_II_316469 | YBR039W | c.895G>T | p.Glu299* | TTF4 |
| chr_IV_135269 | YDL181W | c.91A>T | p.Arg31* | TTD5 |
| chr_X_148659 | YJL141C | c.1732C>T | p.Gln578* | TTG5 |
| chr_XI_239515 | YKL105C | c.3069C>A | p.Cys1023* | TTG5 |
| chr_XII_937991 | YLR410W | c.451G>T | p.Gly151* | TTG5 |
| chr_XIII_409229 | YMR070W | c.76C>T | p.Gln26* | TTA3 |
| chr_XIII_409985 | YMR070W | c.832C>T | p.Gln278* | TTA5 |

**Supplement TableS3 is provided as an excel sheet with the online version of the article.**

### Supplemental Figure Legends

**Figure S1. Quality control of thermoevolved strains after 600 generations.**

(A) 96-well deep plate layout of TT strains selected for phenotype characterization and genome sequencing after 600 generations. Wells included blank media controls for contamination monitoring, with specific arrangements to identify cross-contamination and strains lost during Adaptive Laboratory Evolution (ALE). Color coding: Orange represents S288C strains, Yellow represents BY4741 strains, Grey indicates blank controls, White marks strains lost during ALE, and Black denotes cross-contaminated wells. (B) Microscopy images used to rule out bacterial contamination in TT strains. It was also used to check for cross-well contamination of yeast cells in uninoculated control wells. (C) Ruling out of contamination by mating type PCR. The expected size of PCR products for MATa (BY4741) was 492bp and 369bp for MATα (S288C). (D) Ruling out of contamination by Spotting on SD-Ura plate. To monitor the cross-contamination of S288C(Ura+) with BY4741(ΔUra), TT strains were spotted on the SD-Ura plate. (E) Growth curve of resultant evolved strains at 40°C in YPD medium. TTB1 to TTB11 correspond to evolved thermotolerant strains and BY4741 is the parent strain. The same growth curve of BY4741 is plotted in all the panels for comparison with the evolved strains.

**Figure S2. Genotype characterization from whole genome sequencing reveals routes to thermotolerance.**

(A, B) Drop Dilution assay in YPD agar plate at 30°C and 40°C for selected strains from deletion library with parental unevolved BY4741 as a control. The deletion strain of ecm1 is also replicated in both plates to check for consistency. Growth differences are apparent at the last dilution at 40$^{O}$C. (C) Drop Dilution assay in YPD agar plate at 30°C and 40°C for selected freshly reconstructed deletion strains on the background of unevolved BY4741, along with the parental BY4741 and thermoevolved TTB2 as controls. (D) Competitive fitness of selected strains from deletion library at 40°C showing acquisition of thermotolerance in YPD medium. The data is normalized to parental BY4741 at 40°C. (E) Mutations belonging to similar genes in many TT strains generated from parental BY4741 and S288C were clustered base Jaccard metric. The TT strains associated with genes are shown in the network.

**Figure S3. Thermotolerant strains develop a better tolerance to ER proteotoxicity at higher temperatures**.

(A) Flow cytometry data demonstrates protein translation by using GFP expressed under a constitutive promoter in the evolved strains. (B) MIC assay for TT strains transformed with Plasmid-based protein fC1ing sensors (WTNAT and TS22) in the presence of clonNAT at 34°C in SD-ura with unevolved BY4741 as control. (C) Drop dilution assay of TT strains in the YPD agar plate incubated at 40°C in presence of

Tunicamycin(TM; 1µg/ml), Azetidine-2-carboxylic acid (AZC; 2.5mM) with unevolved BY4741 and ire1Δ as control. (D) Drop dilution assay of TT strains in the YPD agar plate incubated at 40°C in presence of Paraquat (1mM) with unevolved BY4741 as control.

**Figure S4. Rewiring of proteostasis is partially recapitulated by single gene deletions.**

(A) MIC assay of reconstructed deletion strains in increasing concentration of Tunicamycin (TM) in YPD at 30°C. (B) MIC assay of reconstructed deletion strains in increasing concentration of AZC in YPD at 30°C.

**Figure S5. Upregulation of ERAD pathways is prevalent in thermoevolved strains.**

(A) Box plot denoting statistical significance of alterations in ERAD (Endoplasmic Reticulum-Associated Degradation) machinery in TT strains. Significance was calculated using Wilcoxon rank-sum test, with p-value indicated on the graph. (B) Box plot denoting Statistical significance of alterations in ER stress response in TT strains. Significance was calculated using Wilcoxon rank-sum test, with p-value indicated on the graph.

### Supplemental Figures

### Figure S1

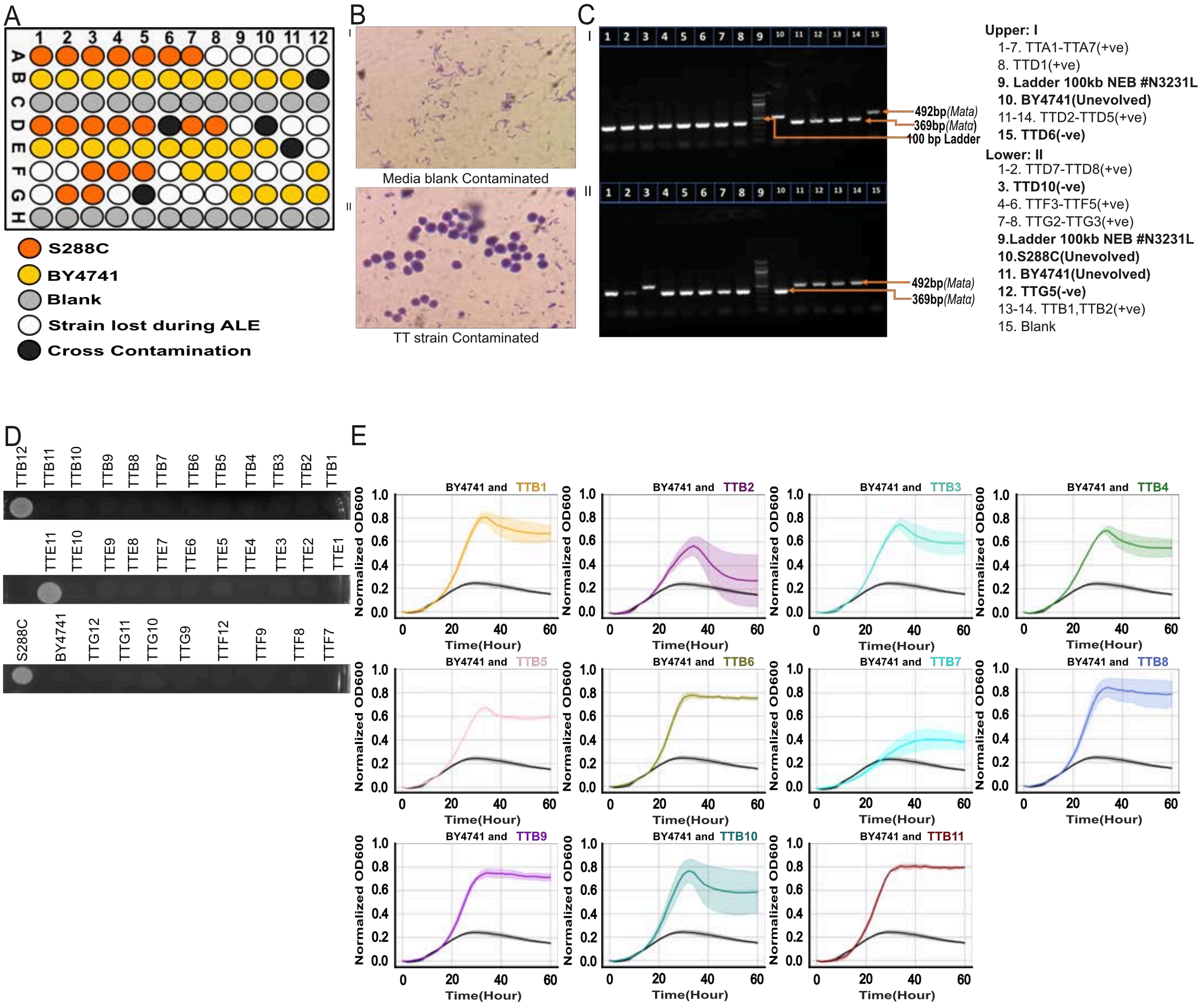
A
1 2 3 4 5 6 7 8 9 10 11 12
A B C D E F G H
S288C
BY4741
Blank
Strain lost during ALE
Cross Contamination
B
I
Media blank Contaminated
II
TT strain Contaminated
C
I
II
492bp(Mata)
369bp(Matα)
100 bp Ladder
Upper: I
1-7. TTA1-TTA7(+ve)
8. TTD1(+ve)
9. Ladder 100kb NEB #N3231L
10. BY4741(Unevolved)
11-14. TTD2-TTD5(+ve)
15. TTD6(-ve)
Lower: II
1-2. TTD7-TTD8(+ve)
3. TTD10(-ve)
4-6. TTF3-TTF5(+ve)
7-8. TTG2-TTG3(+ve)
9.Ladder 100kb NEB #N3231L
10.S288C(Unevolved)
11. BY4741(Unevolved)
12. TTG5(-ve)
13-14. TTB1,TTB2(+ve)
15. Blank
D
TTB12 TTB11 TTB10 TTB9 TTB8 TTB7 TTB6 TTB5 TTB4 TTB3 TTB2 TTB1
TTE11 TTE10 TTE9 TTE8 TTE7 TTE6 TTE5 TTE4 TTE3 TTE2 TTE1
S288C BY4741 TTG12 TTG11 TTG10 TTG9 TTF12 TTF9 TTF8 TTF7
E
BY4741 and TTB1
BY4741 and TTB2
BY4741 and TTB3
BY4741 and TTB4
BY4741 and TTB5
BY4741 and TTB6
BY4741 and TTB7
BY4741 and TTB8
BY4741 and TTB9
BY4741 and TTB10
BY4741 and TTB11
Normalized OD600
Time(Hour)

Figure S2

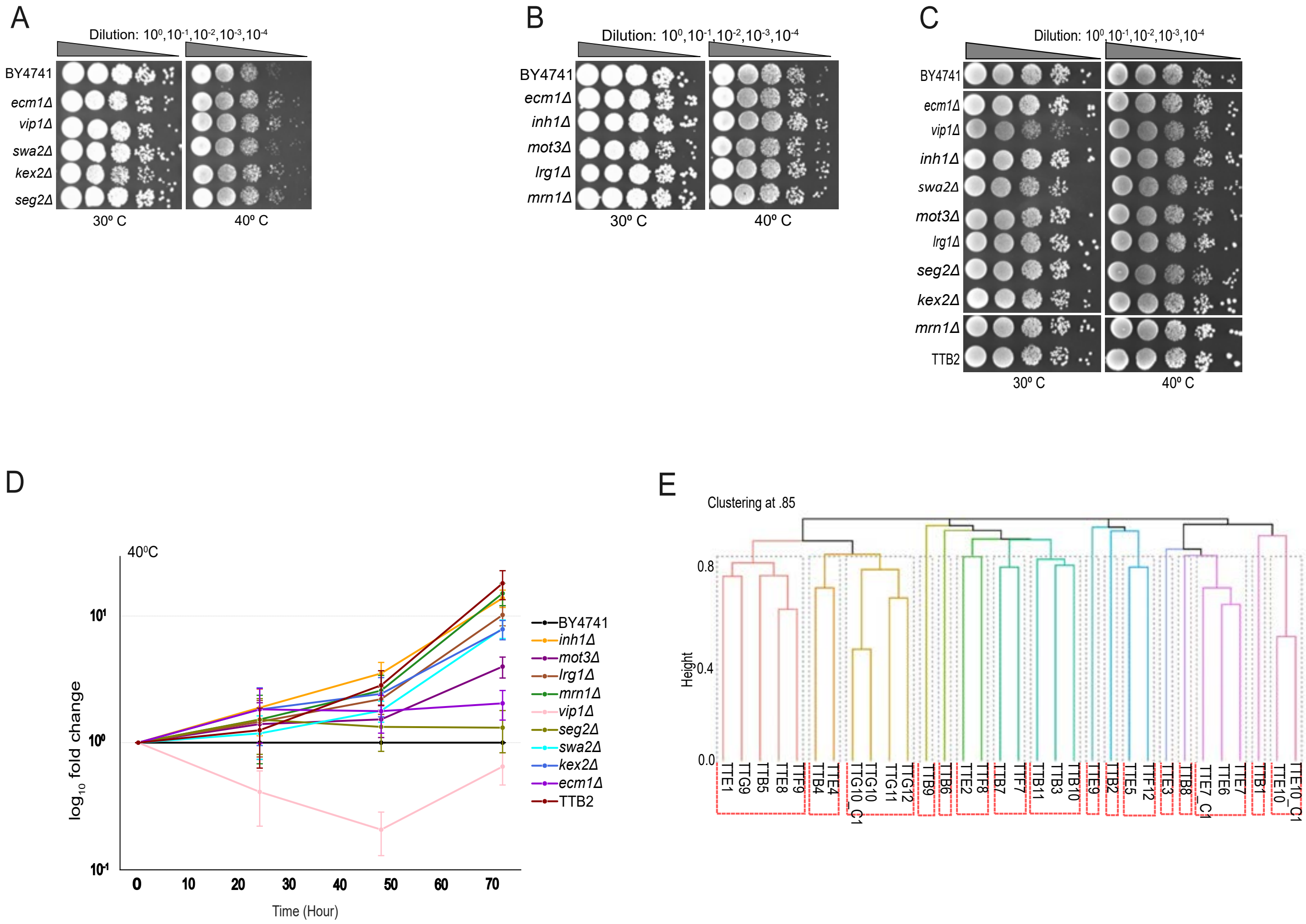
A
Dilution: 10⁰,10⁻¹,10⁻²,10⁻³,10⁻⁴
BY4741
ecm1Δ
vip1Δ
swa2Δ
kex2Δ
seg2Δ
30º C
40º C
B
BY4741
ecm1Δ
inh1Δ
mot3Δ
lrg1Δ
mrn1Δ
C
BY4741
ecm1Δ
vip1Δ
inh1Δ
swa2Δ
mot3Δ
lrg1Δ
seg2Δ
kex2Δ
mrn1Δ
TTB2
D
40ºC
log10 fold change
Time (Hour)
BY4741
inh1Δ
mot3Δ
lrg1Δ
mrn1Δ
vip1Δ
seg2Δ
swa2Δ
kex2Δ
ecm1Δ
TTB2
E
Clustering at .85
Height
TTE1 TTG9 TTB5 TTE8 TTF9 TTB4 TTE4 TTG10_C1 TTG10 TTG11 TTG12 TTB9 TTB6 TTE2 TTF8 TTB7 TTF7 TTB11 TTB3 TTB10 TTE9 TTB2 TTE5 TTF12 TTE3 TTB8 TTE7_C1 TTE6 TTE7 TTB1 TTE10 TTE10_C1

### Figure S3

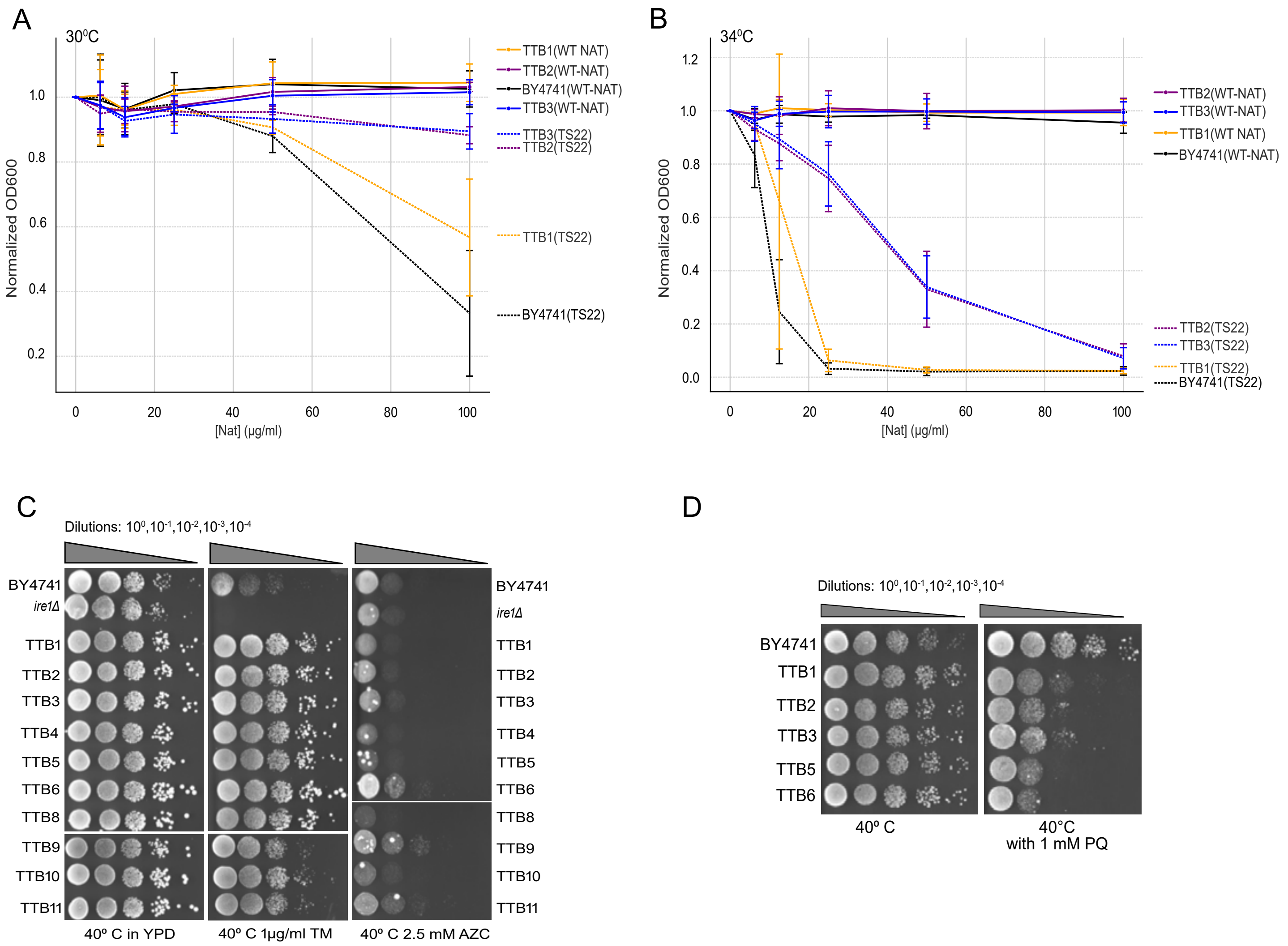
A
30°C
Normalized OD600
[Nat] (µg/ml)
TTB1(WT NAT)
TTB2(WT-NAT)
BY4741(WT-NAT)
TTB3(WT-NAT)
TTB3(TS22)
TTB2(TS22)
TTB1(TS22)
BY4741(TS22)
B
34°C
Normalized OD600
[Nat] (µg/ml)
TTB2(WT-NAT)
TTB3(WT-NAT)
TTB1(WT NAT)
BY4741(WT-NAT)
TTB2(TS22)
TTB3(TS22)
TTB1(TS22)
BY4741(TS22)
C
Dilutions: 10⁰,10⁻¹,10⁻²,10⁻³,10⁻⁴
BY4741
ire1Δ
TTB1
TTB2
TTB3
TTB4
TTB5
TTB6
TTB8
TTB9
TTB10
TTB11
40° C in YPD
40° C 1µg/ml TM
40° C 2.5 mM AZC
D
Dilutions: 10⁰,10⁻¹,10⁻²,10⁻³,10⁻⁴
BY4741
TTB1
TTB2
TTB3
TTB5
TTB6
40° C
40°C with 1 mM PQ

Figure S4

A

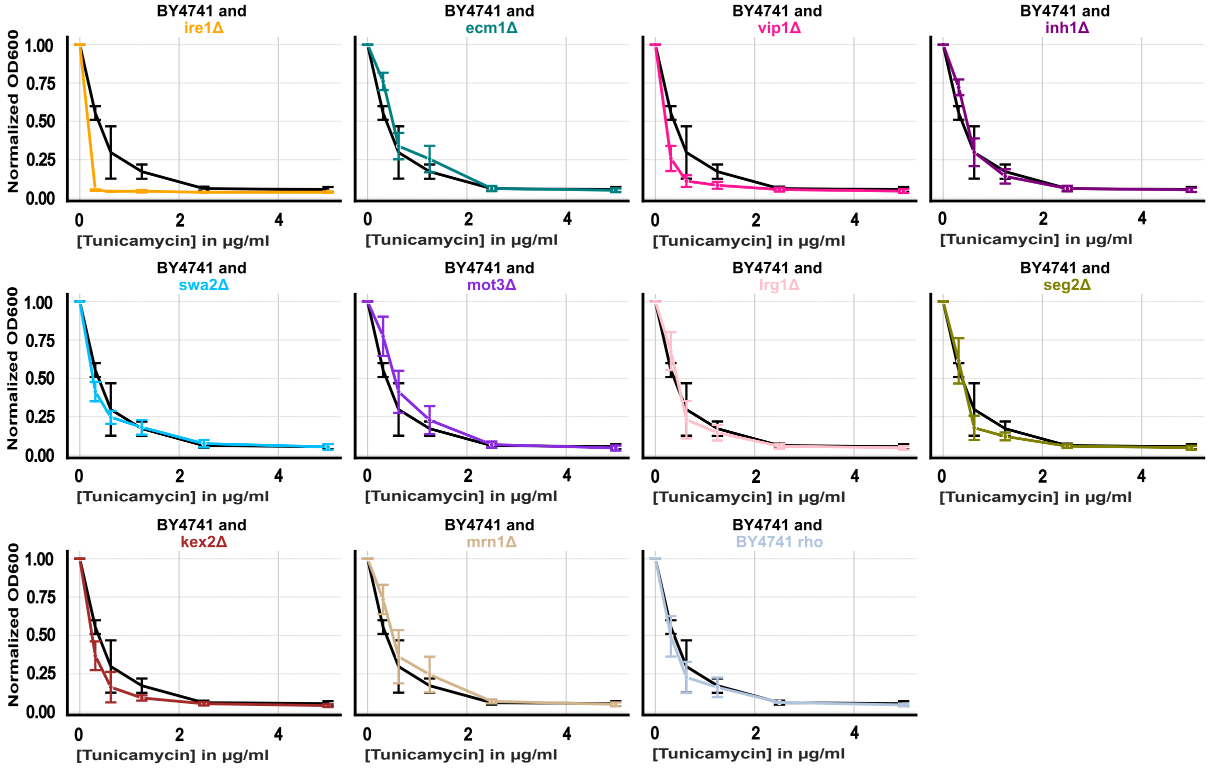
BY4741 and ire1Δ
BY4741 and ecm1Δ
BY4741 and vip1Δ
BY4741 and inh1Δ
BY4741 and swa2Δ
BY4741 and mot3Δ
BY4741 and lrg1Δ
BY4741 and seg2Δ
BY4741 and kex2Δ
BY4741 and mrn1Δ
BY4741 and BY4741 rho
Normalized OD600
1.00
0.75
0.50
0.25
0.00
0
2
4
[Tunicamycin] in µg/ml

B

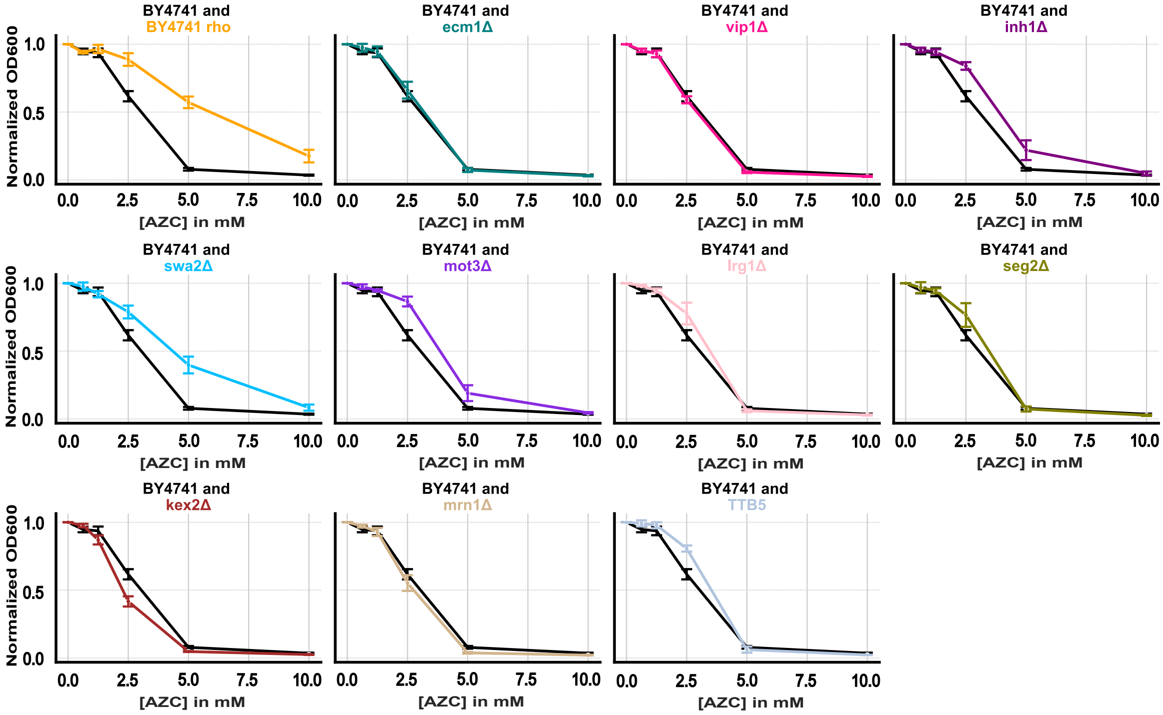
BY4741 and BY4741 rho
BY4741 and ecm1Δ
BY4741 and vip1Δ
BY4741 and inh1Δ
BY4741 and swa2Δ
BY4741 and mot3Δ
BY4741 and lrg1Δ
BY4741 and seg2Δ
BY4741 and kex2Δ
BY4741 and mrn1Δ
BY4741 and TTB5
Normalized OD600
1.0
0.5
0.0
0.0
2.5
5.0
7.5
10.0
[AZC] in mM

### Figure S5

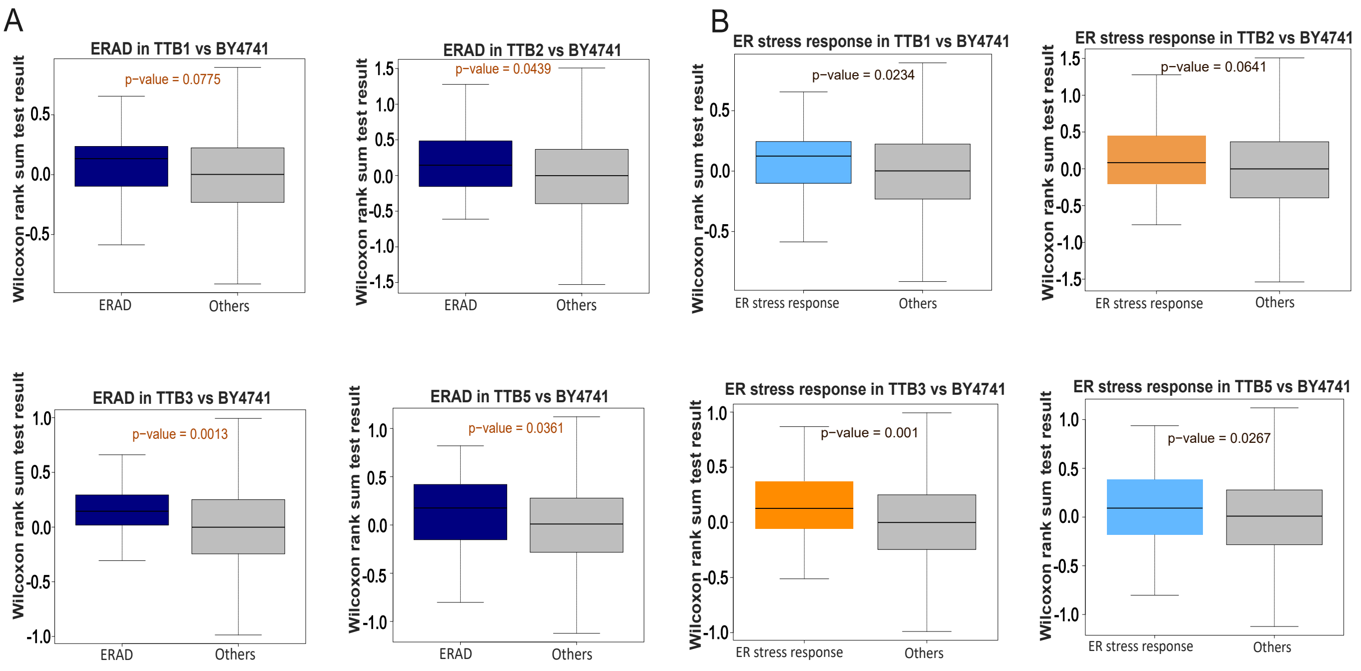
A
ERAD in TTB1 vs BY4741
p−value = 0.0775
ERAD in TTB2 vs BY4741
p−value = 0.0439
ERAD in TTB3 vs BY4741
p−value = 0.0013
ERAD in TTB5 vs BY4741
p−value = 0.0361
ERAD
Others
Wilcoxon rank sum test result
B
ER stress response in TTB1 vs BY4741
p−value = 0.0234
ER stress response in TTB2 vs BY4741
p−value = 0.0641
ER stress response in TTB3 vs BY4741
p−value = 0.001
ER stress response in TTB5 vs BY4741
p−value = 0.0267
ER stress response
Others
